# Extensive collagen deposition by mesenchymal stem cells cultured in 3D self-assembled peptide scaffolds as revealed by nanoplasmonic colorimetric histology

**DOI:** 10.1101/2022.09.15.508197

**Authors:** Christopher J.H. Chong, Vernise J.T. Lim, Mirren Charnley, Julian Ratcliffe, Emily H. Field, Lilith M. Caballero-Aguilar, Chad Johnson, Jacqueline M. Orian, Kristian Caracciolo, Eugeniu Balaur, Brian Abbey, Simon E. Moulton, Katrina J. Binger, Nicholas P. Reynolds

## Abstract

Self-assembling peptides are promising candidates as scaffolds for 3D cell cultures. These hydrogels offer favourable biocompatibility, nanofibrillar structures that mimic native tissues, and the convenient integration of bioactive peptide sequences, such as arginine-glycine-aspartic acid (RGD), which can enable the development of therapeutically valuable cell types.

In the treatment of osteoarthritis (OA) attempts have been made to combine hydrogel scaffolds with mesenchymal stem cells (MSCs) to harness their regenerative potential. This involves the deposition of extracellular matrix (ECM) components like collagen and proteoglycans. Here, we employ the hydrogel-forming peptide Fmoc-diphenylalanine (Fmoc-FF) and incorporate stoichiometric amounts of Fmoc-RGD. We investigate the impact of RGD on nanofibrillar morphologies, hydrogel stability, MSC viability, and the deposition of collagen, proteoglycans, and glycosaminoglycans.

Elevating RGD content enhances cell viability and collagen deposition. However, at higher RGD concentrations, the stability of the hydrogels is compromised. To characterise collagen deposition, we introduce a non-destructive and label-free method using a plasmon-enhanced colorimetric histology technique. This innovation provides a practical means to image collagen without resorting to intricate and destructive sample processing and complex immunohistological staining procedures. This simple approach holds broad potential for routine and label-free quantification of collagen-rich biomaterials, promising widespread applications across various research and clinical settings.

## 1. INTRODUCTION

Osteoarthritis (OA) is a degenerative condition of movable joints which is characterised by degradation of cartilage, joint inflammation and loss of normal joint function. Global Burden of Disease data estimated 595 million prevalent cases of OA worldwide in 2020 (with knees and hips most commonly affected), and with age-standardised prevalence, incidence and years lived with disability (YLDs) rising incrementally by 8-10 % since 1990 (a total case increase of 132.2 % in period to 2020) [1, 2], placing OA as the 10^th^ leading contributor to global YLDs [3]. Current surgical interventions to treat OA are problematic [4], leading to high failure rates and/or poor outcomes [5, 6]. Mesenchymal stem cells (MSCs) are a promising cellular therapeutic to restore cartilage loss that occurs in OA due to their potential to self-replicate, differentiate into chondrocytes while producing the extracellular components of hyaline cartilage (collagen, glycosaminoglycans (GAGs) and proteoglycans (PG)), and can be recruited from accessible tissue reservoirs [7–9]. Direct injection of MSCs into the affected joint is the most common delivery method, but the benefits from this approach are minimal due to poor localisation of cells (engraftment efficiency) [10, 11], limited cell viability [12, 13], inflammation, pain and swelling [14, 15].

Biomaterial scaffolds have been increasingly paired with MSC therapies to overcome some of these repair limitations. A well-suited scaffold for cartilage tissue engineering should protect MSCs when injected into damaged tissue, promote cartilage regeneration through accurate mimicry of chemical and biophysical properties of cartilage extracellular matrix (ECM) [16], and hold a porous architecture which can allow the passage of nutrients and waste products to synovial fluid *in vivo* [17]. Many polymers both natural and synthetic have been explored as MSC scaffolds, and bioinspired self-assembling peptides, such as Fluorenylmethyloxycarbonyl diphenylalanine (Fmoc-FF), combine the advantages of synthetic polymers (purity, ready availability) with a nanofibrillar network structure that mimics the ECM of many tissues [18–24]. Similar nanofibrillar networks have previously been shown to promote cell attachment [25–27] and stem cell differentiation [28, 29].

Fmoc-FF, thanks to its long-range π-π stacking interactions between the Fmoc groups decorating the surface of the nanofibrillar β-sheet rich FF fibrils, rapidly forms hydrogels [24] without the need for any chemical or light-based cross-linking agents that may deleteriously affect cell viability. Fmoc-FF based gels have been shown to readily form stable and transparent hydrogels, that have supported the 3D culture of chondrocytes for up to 7 days [22, 30]. Biochemical functionality can be added to peptide hydrogels by introducing bioinspired functional peptide motifs such as arginine-glycine-aspartic acid (RGD) [31–34]. The concentration of bioactive ligands is a significant consideration in biomaterial design, as an optimal (low) surface concentration of (RGD-containing) fibronectin promotes collagen production in differentiating MSCs (collagen makes up over 80 % of the solid component of cartilage), whilst this effect is diminished or lost at higher concentrations [35].

There have been few reports of hybrid Fmoc-FF, Fmoc-RGD hydrogels finding applications in cartilage tissue engineering, and the RGD motif has successfully been implemented to functionalise more traditional polyethylene glycol (PEG) [36] and hyaluronic acid (HA) hydrogels [37], with their presence contributing towards more robust chondrogenesis, greater sulphonated GAG (sGAG) production and higher chondrogenic gene expression. These hybrid gels have also demonstrated an ability to support MSC viability up to 7 days [33]. However successful examples are rare and translation to the clinic remains elusive.

A significant bottleneck in the successful development of hydrogels for regenerative medicine (especially in delicate non-covalently cross-linked gels like Fmoc-FF) is a lack of non-destructive methods to monitor the processes underpinning ECM deposition *in situ*. Traditional methods such as immunochemistry and histology rely on extensive fixation or cryo-freezing protocols, followed by laborious multi-step staining procedures. All of these processes can destroy scaffolds and alter the presentation of deposited ECM in both research [38] and clinical settings [39, 40]. Molecular techniques, such as qPCR, offer insight on gene expression that might not reflect protein deposition nor spatial distribution. Some efforts have been made to develop improved methods for the *in situ* monitoring of chondrogenesis, including an innovative study by Onofrillo *et. al.* who developed a fluorescent hydrogel to monitor bioscaffolds during neocartilage generation [41]. However, there is still an urgent need to develop simple, preferably label-free and non-destructive methods of assessing the main processes that occur during chondrogenesis (collagen deposition, GAG production, PG production and cell differentiation).

Here, we report a fundamental increase in our understanding of how Fmoc-RGD is incorporated into Fmoc-FF hydrogels, and the development of a new label-free, colorimetric detection of collagen deposition by MSCs encapsulated within the gels. We characterised the biophysical properties of Fmoc-FF hydrogels with stoichiometric amounts of Fmoc-RGD to determine the biodegradation times, nanofibrillar architectures, and mechanical properties of the materials at both the nano- and meso- scale. From these experiments we found that Fmoc-RGD ligands incorporated within Fmoc-FF were able to produce hybrid nanofibrils at low concentrations (1 %), but a phase separation (detrimental to gel formation) occurred at higher concentrations (5 %). To investigate if RGD incorporation affected ECM deposition, we combined existing dimethylmethylene blue (DMMB) and histological assays with an innovative plasmon-enhanced label free colorimetric histology device [42].

The colorimetric histology device is a nanofabricated microscopy slide that exploits the extreme sensitivity of surface plasmon resonance (SPR) to convert subtle changes in the local optical properties of samples adhered to the device surface into a striking colour contrast. In previous work using well-characterised analytes as well as thin films of varying dielectric constant, we showed that bimodal plasmonic devices combined with linearly polarised incident light can be tuned to maximise the sensitivity to certain analytes [43, 44]. Bimodal plasmonic devices also exhibit extreme sensitivity to changes in the local birefringence properties of the sample which manifests as a colour change on the slide. Here we present the first experiments conducted using plasmon-enhanced colorimetric histology to image *in-situ* collagen deposition on the slide surface. By exploiting the colour contrast sensitivity, we extract an estimate of the amount of collagen present on the slide deposited by the MSCs without relying on any staining, labelling or destructive sample preparation.

The colorimetric histology device seamlessly integrates with conventional optical microscopy, aligning with current clinical workflows, as previously demonstrated in cancer studies [42, 45]. In the present context, we envision clinicians potentially utilising this technique to rapidly assess tissue repair by determining the extent of collagen deposition or regeneration in response to implanted scaffolds in vivo. This method requires minimal additional training and equipment, beyond a basic optical microscope.

Our findings not only advance our comprehension of how the inclusion of bioactive elements like RGD in short peptide scaffolds impacts the physical attributes and biocompatibility of 3D cell culture materials but also introduce a novel label-free means of quantifying collagen presence in tissue-engineered scaffolds. The simplicity and speed of this technique are likely to make it an appealing method for assessing collagen deposition across a wide array of biomaterials, both in research and clinical environments. The technique could also be utilised to aid in the design and development of future materials for regenerative medicine.

## 2. MATERIALS and METHODS

### 2.1 Fmoc-FF and Fmoc-FF, Fmoc-RGD Hydrogel Formation

Fmoc-FF hydrogels were formed from a stock solution of Fmoc-FF (Bachem Ltd., Switzerland, purity > 95 %) dissolved in DMSO (100 mg/mL). The stock solution was diluted in Milli-Q^®^ (MQ) ultrapure water to a final concentration of 5.0 mg/mL in glass vials (Samco, United Kingdom). The resulting solution was briefly vortexed and allowed to gelate, then equilibrate over an 8-hour period. Hybrid Fmoc-FF, Fmoc-RGD hydrogels were formed in the same way from stock solutions of Fmoc-FF and Fmoc-RGD (GL Biochem, China, purity > 95 %) (100 mg/mL in DMSO) in appropriate ratios. Three hydrogels were formed at the following ratios 100 % wt Fmoc-FF : 0 % wt Fmoc-RGD, 99 % wt Fmoc-FF : 1 % wt Fmoc-RGD and 95 % wt Fmoc-FF : 5 % wt Fmoc-RGD. For simplicity, for the remainder of this manuscript these gels will be referred to as 0 % RGD, 1 % RGD and 5 % RGD respectively. A schematic representation of the process of fabricating the hydrogels is shown in **figure 1**.

**Figure 1:**
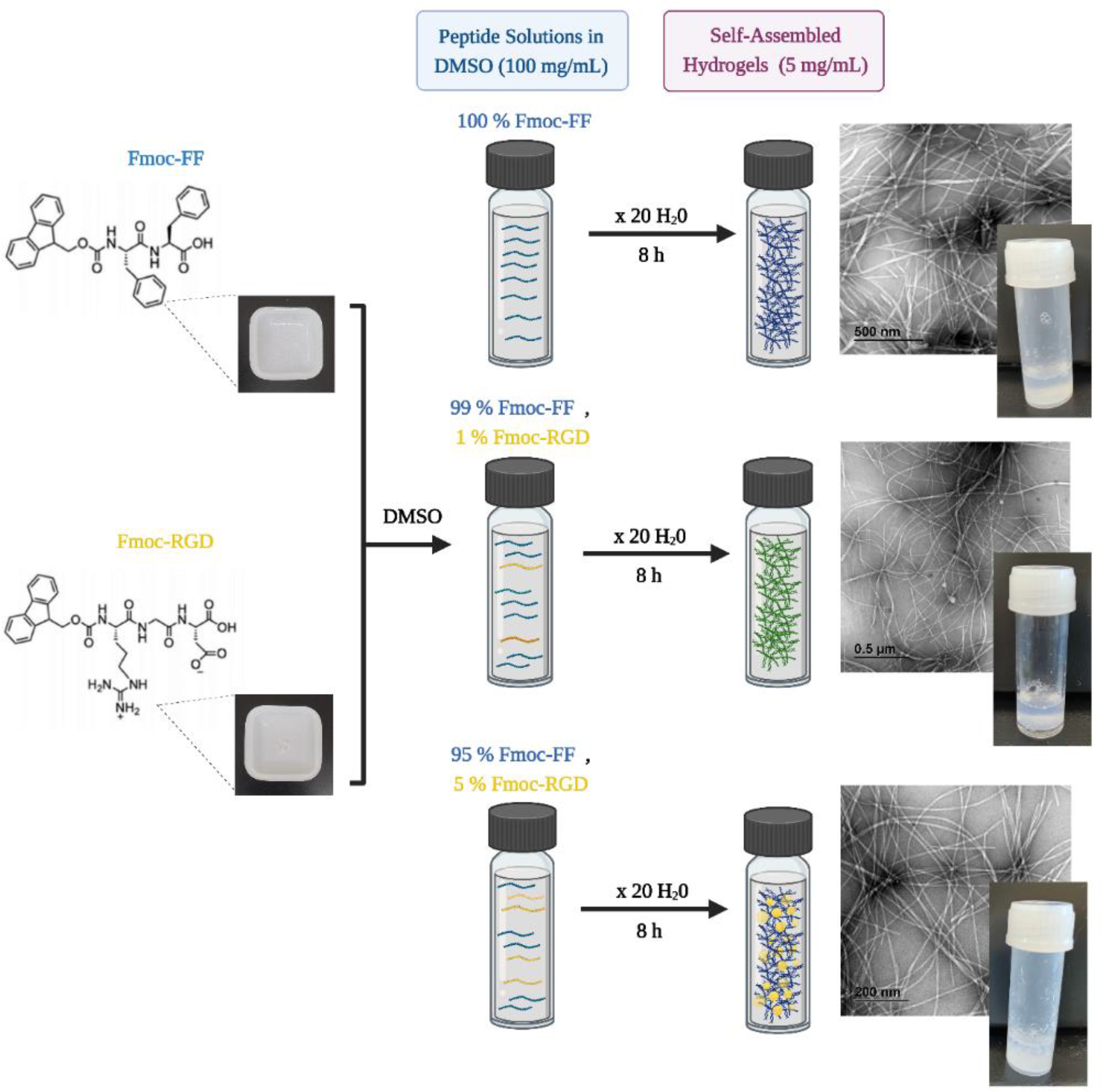
Formation of self-assembled nanofibrillar gels with stoichiometric amounts of RGD ligand. Stoichiometric mixing of Fmoc-FF and Fmoc-RGD in DMSO before dilution into ultrapure water results in the rapid formation of viscoelastic hydrogels. TEM imaging revealed the formation of self-assembled nanofibrillar network arrangements within hydrogels containing 0 %, 1 % and 5 % -RGD. Figure created with Biorender.com.

### 2.2 Degradation Studies

Degradation studies were performed at 37 °C at 5 % CO2. Wash steps on hydrogels were performed every day in either complete hMSC culture medium or phosphate-buffered saline (PBS). 1 mL culture medium or PBS was pipetted at each wash step to hydrogel-containing vials under sterile hood conditions. All media or PBS was then aspirated prior to weighing, and the next wash step being then performed.

### 2.3 Transmission Electron Microscopy (TEM)

TEM analysis was performed using a JEOL JEM-2100 - a 200 kV TEM equipped with two Gatan digital cameras for widefield or high-resolution imaging (a Gatan Orius S200D (2k x 2k) and a Gatan Orius SC1000A (4k x 2.5k)). Approximately 4 µL volumes of gel samples were placed onto glow discharged (Emitech k950x with k350 attachment) carbon coated 300-mesh copper grids (ProSciTech, Queensland, Australia). Excess fluid was drawn off, and the samples were negatively stained with uranyl acetate, wicking away excess after each.

A statistical analysis was conducted through the open-source statistical analysis program, FiberApp (Version 2015, ETH Zurich, Switzerland) [46]. Fibrillar characteristics of interest explored as part of this analysis included fiber width, length, pixel intensity, periodicity of helical fibers, and persistence lengths. Discrete Fourier Transform (DFT) analyses were used to convert the discrete datasets from each set of gel fibers into sequences from which average periodicity measurements were obtained to reflect characteristic distances between intervals of helical arrangements, and widths of protofilaments within helical arrangements.

### 2.4 Rheology

Rheology was performed using an Anton Paar MCR-302 rheometer (Anton Paar GmbH, Graz, Austria). All data was collected with Anton Paar RheoCompass^TM^ software (Anton Paar GmbH, Graz, Austria) [47], and processed with OriginPro (Version 2020, OriginLab Corporation, Northampton, MA, USA) [48] to determine the storage modulus (**G’**) and loss modulus (**G’’**) of the gels. The measurements were performed using a parallel plate geometry (10 mm plate) with 0.05 N force at room temperature, using a constant angular frequency of 10 rad s^-1^.

Oscillatory measurements were performed either at constant frequency (1 Hz) over 300 seconds, or in a frequency sweep (0.1-100 Hz). All samples were run in triplicate and the data plotted as a mean value ± one standard deviation. Experiments were conducted to understand the viscoelastic properties of each hydrogel type with relation to changes in shear strain.

### 2.5 Small Angle X-ray Scattering (SAXS)

Small Angle X-ray Scattering (SAXS) experiments were performed at room temperature on the SAXS / WAXS (Wide-Angle X-ray Scattering) beamline at the Australian Synchrotron. The three peptide assemblies (0 % RGD, 1 % RGD and 5 % RGD) were prepared in dilute solutions to prevent gelation (0.5 mg/mL) in Milli-Q^®^ (MQ) water and left to equilibrate for 24 hours at room temperature. 100 µL of each assembly was loaded into a 96-well plate held on a robotically controlled x-y stage and transferred to the beamline via a quartz capillary connected to a syringe pump. The experiments used a beam wavelength of *λ* = 1.03320 Å (12.0 keV) with dimensions of 300 *µ*m × 200 *µ*m and a typical flux of 1.2 × 10^13^ photons per second. 2D diffraction images were collected on a PILATUS 1M detector at *q*-ranges between 0.02 and 0.25 Å^−1^. Spectra were recorded under flow (0.15 mL/min) to prevent x-ray damage from the beam. Multiples of approximately 15 spectra were recorded for each time point (exposure time = 1 second) and averaged spectra are shown after background subtraction against MQ water in the same capillary.

### 2.6 Cell Culture

Human bone-marrow derived mesenchymal stem cells (hMSCs) were obtained from Lonza (PT-2501, Basel, Switzerland). A complete hMSC culture medium was formed from Dulbecco’s Modified Eagles’ Medium (DMEM - low glucose) (Sigma-Aldrich Ltd.), foetal bovine serum (FBS) (10 %, 12003C-500 mL, Sigma-Aldrich Ltd.), HEPES (15 mM, purity ≥ 99.5 %, H4034-500 g, Sigma-Aldrich Ltd.), L-Glutamine (2 mM, 59202C, Sigma-Aldrich Ltd.), antibiotics Penicillin-Streptomycin (1X, P4333-100 mL, Sigma-Aldrich Ltd.), Epidermal Growth Factor (20 ng/mL, E9644-0.2 mg, Sigma-Aldrich Ltd.), and Fibroblast Growth Factor (1 ng/mL F3685-25 µG, Sigma-Aldrich Ltd.). Cells were cultured in the hMSC culture medium in T-75 or T-175 tissue cell culture flasks (Greiner CELLSTAR^®^, Sigma-Aldrich Ltd.) and incubated at 37 ⁰C in an atmosphere of humidified 5 % CO2 / 95 % air. The media was changed every second day, until such time as the cells reached 90 % confluency and were ready for passage. Cells were used between passages 1-14 (P1-P14). Cell counting was performed with a haemocytometer slide (4 x 16 well grid) and the addition of Trypan Blue cell stain solution (0.4 %, 15250061, Gibco, New York, USA), with samples viewed under a Nikon Eclipse TS100 inverted microscope (Nikon, Tokyo, Japan).

### 2.7 Live-Dead Cell Assay

A Live-Dead Cell Staining Kit (04511-1KT-F, Sigma-Aldrich Ltd.) was used for simultaneous fluorescence staining of viable and dead cells. The stain kit contains Calcein-AM (which is converted to a highly fluorescent molecule by metabolically active cells (green)) and Propidium Iodide (PI) (which stains the DNA of dead or dying cells (red)). A modified staining protocol was used to account for the 3D matrix, with MSC laden hydrogels (250,000 cells per mL) evaluated after 1, 3, 6 days in culture – involving a wash twice in PBS (15 minutes) before incubation of cells within a stain solution of Calcein-AM and PI, in PBS for 60 minutes. Samples were then washed twice more in PBS (15 minutes) before imaging on a dual channel fluorescent microscope (Olympus IX81, Olympus, Tokyo, Japan).

### 2.8 Fluorescence Microscopy

Imaging was performed and captured with the Olympus IX81 fluorescence microscope (20X and 40X magnification) and Olympus Dimension Cellsens software (Olympus, Tokyo, Japan). Fluorescein / Green fluorescent protein / Alexa Fluor 488 (FITC / GFP / AF488), and Tetramethylrhodamine-isothiocyanate / Alexa Fluor 568 (TRITC / AF568) were used as dual band filter channels to capture fluorescence from live and dead cells respectively. A Live-Dead Cell Staining Kit (04511-1KT-F, Sigma-Aldrich Ltd.) was utilised for simultaneous fluorescence staining of viable and dead cells, with excitation of both stains (Calcein-AM and Propidium Iodide) induced at 490 nm, and emission wavelengths captured at 518 nm (FITC), and 580 nm (TRITC) respectively.

### 2.9 Resazurin-based Cell Assay

0 % RGD, 1 % RGD and 5 % RGD gels were formed to volumes of 500 μL in black 96-well MicroWell plates (Nunc, Roskilde, Denmark), and incubated for 24 hours. Gels were then washed extensively with hMSC media and seeded with 72,000 MSCs per well for a further 24 hours. To determine cell viability, gels were washed twice with PBS before incubation with the cell permeable, resazurin-based Presto Blue^TM^ assay solution (Sigma-Aldrich Ltd.) as per the manufacturer’s recommendations. Resazurin is reduced to the highly fluorescent molecular resorufin via cellular aerobic respiration of metabolically active cell populations and can then be detected through measurement of fluorescence. Fluorescence was determined by a CLARIOstar Microplate Reader (BMG Labtech, Ortenberg, Germany) with excitation at 535 nm and emission 615 nm. All samples were run in triplicate and the data plotted as a mean value

± one standard deviation. Statistical analysis was performed in Graphpad Prism (version 10.0.2 for Windows, GraphPad Software, Boston, Massachusetts USA, www.graphpad.com) [49] using a one-way ANOVA with Tukey comparison.

### 2.10 ECM deposition experiments

To quantify the amounts of collagen, sGAGs and proteoglycans (PGs) deposited by the MSCs in the different hydrogels, long-term cell culture experiments were performed. 0 % RGD, 1 % RGD hydrogels were formed in screw cap Eppendorf vials as per section 2.1. Pure (100 %) collagen gels were formed from dissolution of PureCol® EZ gel neutralised type I collagen solution (bovine, advanced biomatrix) into Milli-Q^®^ (MQ) ultrapure water to a final concentration of 2.5 mg/mL. MSCs were seeded at 250,000 cells per mL and cultured for 3 days in the MSC expansion media detailed in section 2.6. After this initial cell expansion, the media was changed into either chondrogenic promoting media (chondro media: DMEM high glucose media (11-965-084, Gibco) + Insulin Transferrin Selenium (ITS) 0.01X (I3146- 5mL, Sigma-Aldrich Ltd.) + 100 nM water-soluble Dexamethasone (D2915-100MG, Sigma-Aldrich Ltd.) + 50 µg/mL Ascorbic Acid-2 Phosphate (A8960-5G, Sigma-Aldrich Ltd.) + 10 ng/mL TGF-β (SRP3171, Sigma-Aldrich Ltd.) + 10 ng/mL BMP-6 (SRP3017, Sigma-Aldrich Ltd.) or non-chondrogenic promoting media (chondro media without the addition of TGF-β or BMP-6 (basal media)) for 21 days, with media changes occurring every 2 or 3 days.

### 2.11 Colorimetric Histology

The planar plasmonic devices used for the current study were fabricated using displacement Talbot lithography (Phabler 100C, Eulitha AG). The plasmonically active layer is fabricated in a 150 nm thick silver (Ag) film, incorporating circular nanoapertures (180 nm diameter) on a rectangular pattern with a 400 nm period in one direction (corresponding to ‘s-polarised’ illumination) and a 500 nm period in the orthogonal direction (corresponding to ‘p-polarised’ illumination). The two different periodicities mean that quite distinct plasmon-resonance mode structures can be excited simply by rotating the incident polarisation vector of the light between orthogonal directions via means of a linear polariser inserted just below the eyepiece of the microscope. To protect the surface from the surrounding environment, a very thin layer (10 ± 1 nm) of SiO2 was deposited as the final step after fabrication (see e.g. Balaur *et. al.* [42] for further details of the nanofabrication). Small aliquots of each of the gel-cell constructs were pipetted and seeded onto the plasmonic devices. Samples were imaged on the Nikon Ti2 inverted microscope (Nikon, Tokyo, Japan), and two images recorded with either s-polarised or p-polarised incident light. Rotating the incident polarisation produced two very distinct colour contrast images. Any birefringence of the sample induces an additional rotation of the polarisation vector which can be qualitatively characterised as a change in colour from s to p polarisation (or vice versa). To investigate the collagen content of our samples, areas within cell-seeded (0 % RGD, 1 % RGD) gels were compared with pure collagen gels and gels without cells (0 % RGD, 1 % RGD).

An estimate of the collagen deposition observed via colorimetric histology was performed in ImageJ [50]. Colour images were converted to HSB (Hue, Saturation and Brightness) stack images, and a threshold set (206/255) in the brightness channel for the pure collagen gel (s-polarised) eliminating all background signal not arising due to collagen birefringence. The same thresholding was applied to all images, and the ratio of black to white pixels in the Hue channel was determined for 3 independent images for each condition (chondrogenic media 0 % and 1 % RGD and basal media 0 % and 1 % RGD). Statistical analysis was performed in Graphpad Prism [49] using a one-way ANOVA with Tukey comparison.

### 2.12 Histology

A Safranin O staining protocol for cartilage was utilised to assess PG protein expression on 0 % RGD, and 1 % RGD. All Fmoc gels were exposed to either chondrogenic or basal media, and samples were collected at days 0 and 21. A manual 10-step sequence of handling was used to fix the samples – first with paraformaldehyde (4 % w/v) (2 hours), 70 % ethanol (10 minutes), 100% ethanol (twice for 30 minutes, then once for 45 minutes), then xylene (twice for 30 minutes, then once for 45 minutes) (total exposure time: 5 hours, 40 minutes). Samples were embedded directly using the LEICA EG 1150H paraffin embedder (Wetzlar, Germany) within paraffin moulds and cooled on the LEICA EG 1150C cold plate. Gel-cell samples, were placed in the base of the mould and layers of paraffin were dispensed until the chamber was full. Gel-cell samples were generally sectioned into fragments randomly prior to embedding to maximise exposed surface area to wax. A further layer coating of paraffin was dispensed, and the samples were then quickly relocated to a cooling counter for 10 minutes. Samples were soaked in paraffin wax for 3 days, and then re-embedded in a wax refill.

Paraffinised samples were sectioned at 7 µm thickness with a LEICA RM2235 rotary microtome (Wetzlar, Germany) for Safranin O staining. These sectioned samples were briefly transferred to a water bath (42 °C), before loading onto microscope slides (ThermoScientific Menzel Gläser Superfrost^®^ Plus), samples were allowed to anneal and dry in room air overnight.

A modified Safranin O protocol was adapted from Tran et al. (2000) [51]. 0.1 % (w/v) Safranin O stain solution was formed from dissolution of Safranin O (colour index 50240) in MQ. Safranin O staining of samples required deparaffinisation (dewaxing) of samples and hydration of slides through a series of exposure washes to xylene (2 x 2 minutes), pure ethanol (2 x 1 minute) and tap water (5 minutes). Samples were then stained in 0.1 % Safranin O solution for 5 minutes. Samples were finally dehydrated and cleared with 70 % ethanol (10 dips), pure ethanol (2 x 10 dips) and xylene (2 x 1 minute), and then a coverslip was applied on the surface of the section.

Images of sample slides were captured with the LEICA Aperio AT2 digital pathology system (Wetzlar, Germany), and ImageScope (Version 12.4.6, LEICA, Wetzlar, Germany) [52]. Using the batch scanner and image stitch functions made available by the Aperio and the Zeiss Axioscan 7 (Oberkochen, Germany), whole slide images were obtained for any remaining viable slides. Images of each sample were captured using the 20x lens. The HALO^®^ Image Analysis Platform (Version 3.2, Indica Labs, New Mexico, USA) [53] was then utilised to quantitatively classify regions of Safranin O-stained tissue features on each slide (correlating with PG expression), to determine proportionate PG expression relative to other slide features (glass surface area, particulates etc.) across selected regions of 500 µm^2^. Statistical analysis was performed in Graphpad Prism [49] using a one-way ANOVA with Tukey comparison.

### 2.13 Dimethylmethylene Blue (DMMB) Assay

DMMB (C18H22ClN3S, Sigma-Aldrich Ltd.) is a dye that stains sGAGs, when binding occurs between the dye and these polysaccharides. All gels were prepared in the same manner, briefly: cell laden gels were digested in a papain solution (papain (250 μg/mL, Sigma-Aldrich Ltd.) + 0.01 M cysteine + 0.2 M sodium phosphate + 0.01 M EDTA) overnight at 65°C in a heating block. Following digestion, 100 µL of sample was transferred to a 96 well plate and 100 µL DMMB solution was added (0.348 mM DMMB (Sigma-Aldrich Ltd.) + 34.78 mM sodium chloride (Sigma-Aldrich Ltd.) + 66.09 mM glycine (Sigma-Aldrich Ltd.) + 0.1 M acetic acid (Sigma-Aldrich Ltd.). Absorbance was measured immediately after adding DMMB dye using the CLARIOstar^®^ plate reader (BMG Labtech, Offenburg, Germany, 595 nm). Background absorbance readings were obtained for dye solution alone and subtracted from obtained sample read absorbances.

## 3. RESULTS & DISCUSSION

### 3.1 Hybrid Fmoc-FF, Fmoc-RGD hydrogels degrade more rapidly than pure Fmoc-FF hydrogels

Degradation studies were completed to compare the stability of the 0 % RGD, 1 % RGD and 5 % RGD containing gels. All 3 hydrogels formed self-supporting hydrogels in pure water (**figure 2a**) and their structure was maintained after washing in cell culture media. We observed rapid mass loss within the hybrid (1 % and 5 % RGD) gels in response to cell culture media washes at a rate up to 1.5 times greater than pure Fmoc-FF gels (**figure 2b**). Overall, the 5 % RGD gels completely degraded more rapidly than the 1 % gels (**figure 2b**). In DMEM, the 5 % RGD gels degraded to half of their initial mass within two days (**figure 2b**), whereas 0 % RGD and 1 % RGD gels reached this same degradation point within 32 and 36 days respectively (**figure S1**). These results suggested that 5 % RGD gels were unsuitable for long-term 3D cell culture applications. Similar results were seen for gels washed in PBS (**figure S2a**), and for a larger volume of gels (**figure S2b-c**).

**Figure 2:**
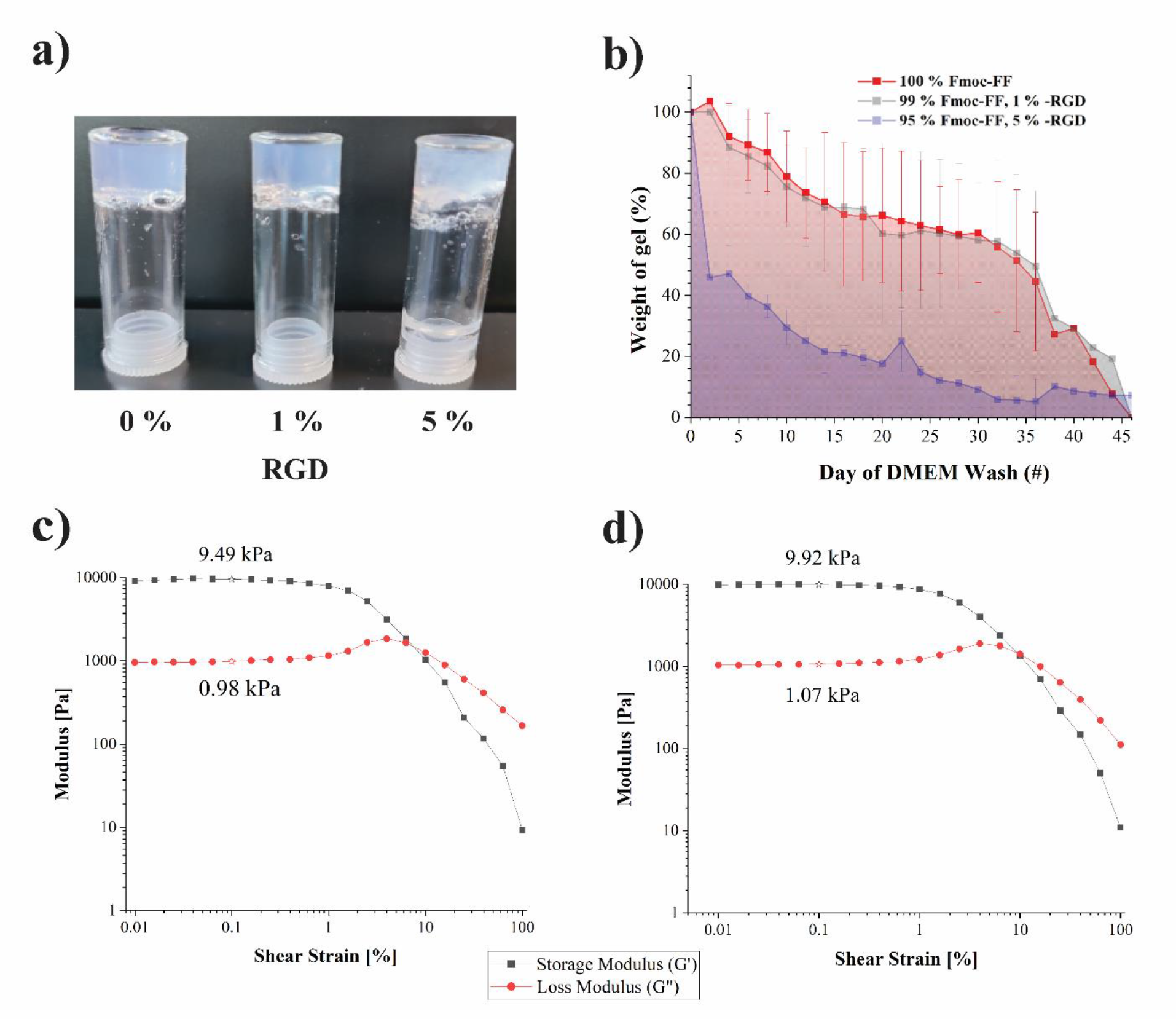
Increasing RGD concentration reduces the stability of the viscoelastic gels. All concentrations of RGD tested form of **a)** self-supporting hydrogels. Increasing RGD concentration results in rapid gel degradation, as shown by **b)** mass-loss assays. Rheology reveals that **c)** 0 % RGD and **d)** 1 % RGD result in viscoelastic hydrogels with storage moduli (G’) of approximately 10 kPa. Storage and loss moduli at 0.1 % shear strain marked on plots for comparison **(c and d)**.

The macroscale mechanical properties of the gels were investigated by rheology, and in the linear viscoelastic region (LVR) the storage moduli (**G’**) of 0 % RGD and 1 % RGD gels were observed to be larger than their respective loss moduli (**G”**) (**figure 2c-d**), indicating the gels are a viscoelastic solid. Storage moduli for both gels in the LVR were between 9.5-10 kPa, which is typical for soft physically crosslinked gels. The respective rheological breaking points of the 0 % and 1 % RGD gels, as indicated by crossover between storage and loss moduli curves occurred at 6.31 % and 10 % shear strain respectively, demonstrating approximately comparable gel stabilities. The 5 % RGD gels were too heterogenous to obtain reliable viscoelastic modulus data, therefore are not included. In addition, rheological analysis revealed that these gels were stable under oscillatory strain up until 60 Hz and 48 Hz respectively, where they underwent a viscoelastic solid-liquid state change (**figure S3**).

### 3.2 All gels are formed from nanofibrillar networks, but TEM reveals subtle differences in their morphologies

To further investigate the nanoscale structure of the self-assembling peptides we turned to transmission electron microscopy (TEM). Negative stain TEM imaging (**figure 3**) revealed a network of nanofibrils within all three hydrogels. Closer inspection highlighted that all three nanofibrillar varieties (0 %, 1 % and 5 % RGD) displayed polymorphic behaviour with mixtures of linear nanofibrils interspersed with those possessing a helical or twisted ribbon structure [54] (**figures 3a-d and S4)**. These regular twists in fibres and flat ribbons have been previously described in TEM imaging of Fmoc-FF / -RGD hydrogels [55].

**Figure 3:**
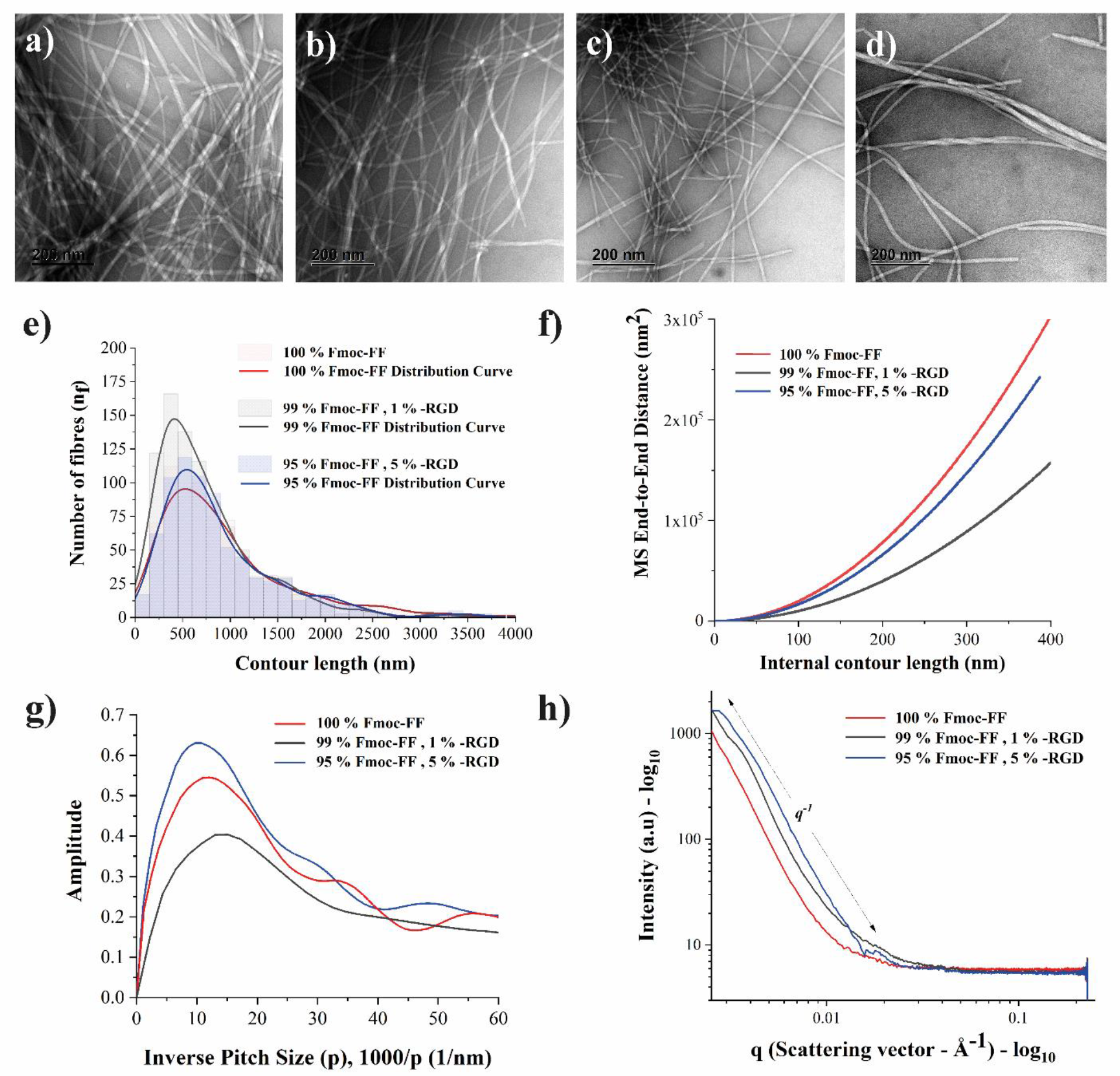
TEM reveals subtle differences in nanofibrillar structure upon the introduction of RGD. Biophysical analysis of the self-assembled nanofibrils reveals distinct nanoscale differences between the gels. **a-d)** TEM images of: **a)** 0 % RGD **b)** 1 % RGD and **c)** 5 % RGD hydrogels. **d)** An example of the periodically twisted ribbon-like nanofibrils seen in all 3 systems. **e-g)** Statistical analysis of the TEM images of the nanofibrils - all analysis performed on more than 500 nanofibrils: **e)** Nanofibril contour length distribution, **f)** Mean-square end-to-end distance of the persistence length (λMSED), **g)** Direct Fourier Transform (DFT) analysis of the pitch of the twisted ribbon-like nanofibrils observed, **h)** Small-angle X-ray scattering (SAXS) analysis of the three nanofibrillar samples at non-gelling concentrations.

Single fibril analysis of the TEM images revealed subtle differences in the nanoscale architecture of the three nanofibrillar networks studied. Quantification of the contour length distributions of the fibrils in the 1 % RGD gels showed that the fibrils tended to be shorter, with the mean of the 0 % RGD, 1 % RGD and 5 % RGD gels measured at 529, 418 and 544 nm respectively (**figure 3e**). The mean square end-to-end distance (MSED) persistence length (λMSED) of the fibers (**figure 3f**) within the 1 % RGD gels was less than the other two gels studied (λMSED = 3.51 µm, 1.77 µm and 2.38 µm for 0 % RGD, 1 % RGD and 5 % RGD gels respectively). Discrete Fourier Transform (DFT) analysis (**figure 3g**) of the periodicity of the fibers with a clear twisted ribbon architecture once again revealed the 1 % RGD gels to be distinct from the 0 % RGD and 5 % RGD gels. The peak-to-peak periodicity of the 1 % RGD gels was just 70.7 nm (from **figure 3g -** 1000/14.15 nm^-1^), whilst this value was 87.4 nm and 93.7 nm for the 0 % RGD gels and the 5 % RGD gels. These findings collectively highlight that there is a distinct difference in the biophysical properties of the nanofibrils found in 1 % RGD gels compared to the other two systems.

The distinct nature of the fibrils in the 1 % RGD samples can be explained in terms of the molecular incorporation of Fmoc-RGD into the self-assembling fibrils. At low concentrations of Fmoc-RGD, these ligands co-assemble with the Fmoc-FF peptides and produce nanofibrils (**figure 3b-d)** that are distinct from the pure Fmoc-FF populations (0 % RGD, **figure 3a**). Once the concentration of Fmoc-RGD ligands increases past a threshold (between >1 % and 5 % RGD), insertion of Fmoc-RGD into the assembling nanofibrils is lost, and instead the Fmoc-RGD ligands non-specifically aggregate into unstructured colloidal assemblies, leaving behind a network of nanofibrils similar to that of the pure Fmoc-FF gels (hence the fibrils in the 5 % RGD samples more closely resemble the fibrils in the 0 % RGD samples than the fibrils in the 1 % RGD samples). The presence of the hypothesised Fmoc-RGD colloids now reduces the long-range order of the nanofibrils formed, preventing the stable gelation of the 5 % RGD hydrogels. Similar phenomena have been observed when adding polysaccharides to Fmoc derived self-assembling peptides, where the gels can incorporate low concentrations of polysaccharide and behave similarly to the pure peptide gels, but at higher concentrations the non-assembling behaviour of the polysaccharide begins to dominate [56].

Support for the presence of colloidal aggregates of Fmoc-RGD at 5 % comes from the SAXS data (**figure 3h**) which shows that the 0 % RGD and the 1 % RGD gels have form factors with a slope approximating q^-1^ (scattering vector) indicative of long rod-like structure (nanofibrillar) formation [57–59]. For these two assemblies, the intensity of scattering continues to increase exponentially past the limit of these plots (∼ 0.002 Å^-1^) due to the contour length of the fibrils being much larger than can be detected at qmin (minimum scattering vector). The 5 % RGD gels, whilst still being dominated by this rod-like form factor (still with an abundance of nanofibrils), start to plateau at low q values. This plateau is likely indicative of the beginning of a Guinier region and the presence of smaller amorphous aggregates of Fmoc-RGD that do not exceed the maximum observable dimensions of the instrument [58]. However, it should be noted that the size of this apparent Guinier region was too small to perform a reasonable analysis of the radius of gyration (Rg) of these amorphous aggregates, this is likely due to the heterogeneous nature (fibrils plus aggregates) of the 5 % RGD gels.

### 3.3 RGD concentration affects MSC viability in hydrogels over time

We next examined the viability of MSCs cultured within the three gels. Live-dead assays (**figure 4a-c**) revealed that most cells cultured in each of the gels remained viable after 24 hours. As previously observed, this fits with previous suggestions that Fmoc-FF based hydrogels are suitable for MSC [60–62] and chondrocyte [22, 63] cell cultures. Imaging of the 5 % RGD gels showed non-continuous gels broken into small fragments (**figure 4c**), that whilst still able to support viable MSCs, these did not remain in a self-supporting hydrogel structure. This again highlights the unsuitability of the 5 % RGD system for long term cell culture applications. A resazurin based metabolic activity (Presto Blue^TM^) assay showed statistically similar levels of metabolic activity between 0 % RGD and 1 % RGD gels over 6 days. 1 % RGD gels only demonstrated slightly greater fluorescence (7-fold increase - **figure 4d**) after 6 days when compared to 0 % RGD gels (4-fold increase - **figure 4d**). In contrast, 5 % RGD hydrogels displayed a high amount of metabolic activity after one day, which is likely due to the high concentration of RGD ligands which promote cell attachment and viability. However, after three days, the metabolic activity dropped; likely due to premature degradation of the 5 % RGD gels. With exposure to repeated media washes, much of the semi-degraded gel was washed away, along with the encapsulated cells (**figure 4d**). This loss of cells due to repeated washes and increased gel degradation explains the apparent drop in cell number after one day in the 5 % RGD gels.

**Figure 4:**
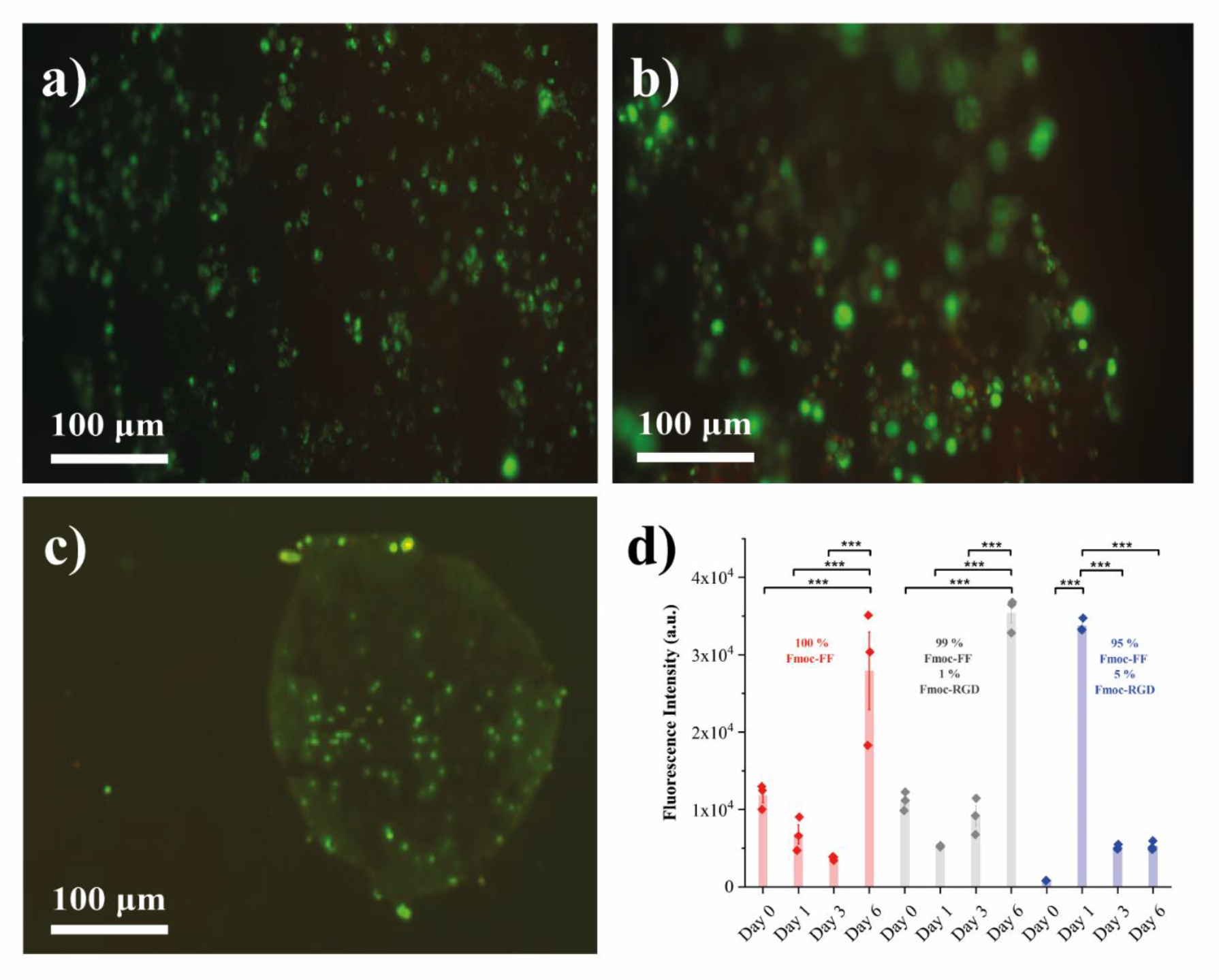
**All gels support MSC growth, but the instability of the 5 % RGD gels makes them unsuitable for long-term culture**. Live-dead assays of mesenchymal stem cells (MSCs) seeded upon **(a)** 0 % RGD **(b)** 1 % RGD and **(c)** 5 % RGD hydrogels after 24 hours culture. Cells were stained with calcein AM (green) and propidium iodide (red). Images captured at 20x magnification on Olympus IX81 fluorescence microscope. **(d)** Resazurin based metabolic activity assay (PrestoBlue^TM^) on the 3 gels studies, error bars show ± 1 standard deviation. *** = P <0.001.

### 3.4 The presence of RGD in the scaffolds resulted in no significant increases in proteoglycan or glycosaminoglycan content

The results above suggest that Fmoc-RGD can only be incorporated into the Fmoc-FF gels at low concentrations. We next performed experiments to determine if the addition of RGD at 1 % promotes the deposition of ECM proteins (collagen, PGs and sGAGs). Pure Fmoc-FF hydrogels and hydrogels containing 1 % RGD were seeded with MSCs (250,000 cells per mL) for 3 days in MSC expansion media. After this time the gels were either transferred to media that promotes chondrogenic differentiation (chondro media) or a control media that does not contain chondrogenesis promoting growth factors (basal media). Additionally negative control samples were taken before the 21-day chondrogenesis commenced (termed D0 samples). All of these samples were then assayed for PG, sGAG and collagen deposition.

PG content was determined using paraffin-based histology, Safranin O dye staining (a histochemical stain that provides contrast to tissue sections, particularly for PG content) (**figure 5, S5**) and quantified with the HALO^®^ Image Analysis Platform [53]. The delicate nature of the non-covalently cross-linked hydrogels made this histological preparation challenging, occasionally leading to sample loss - consequently, we present this data as the mean of all successful technical replicates (across three biological replicates).

**Figure 5:**
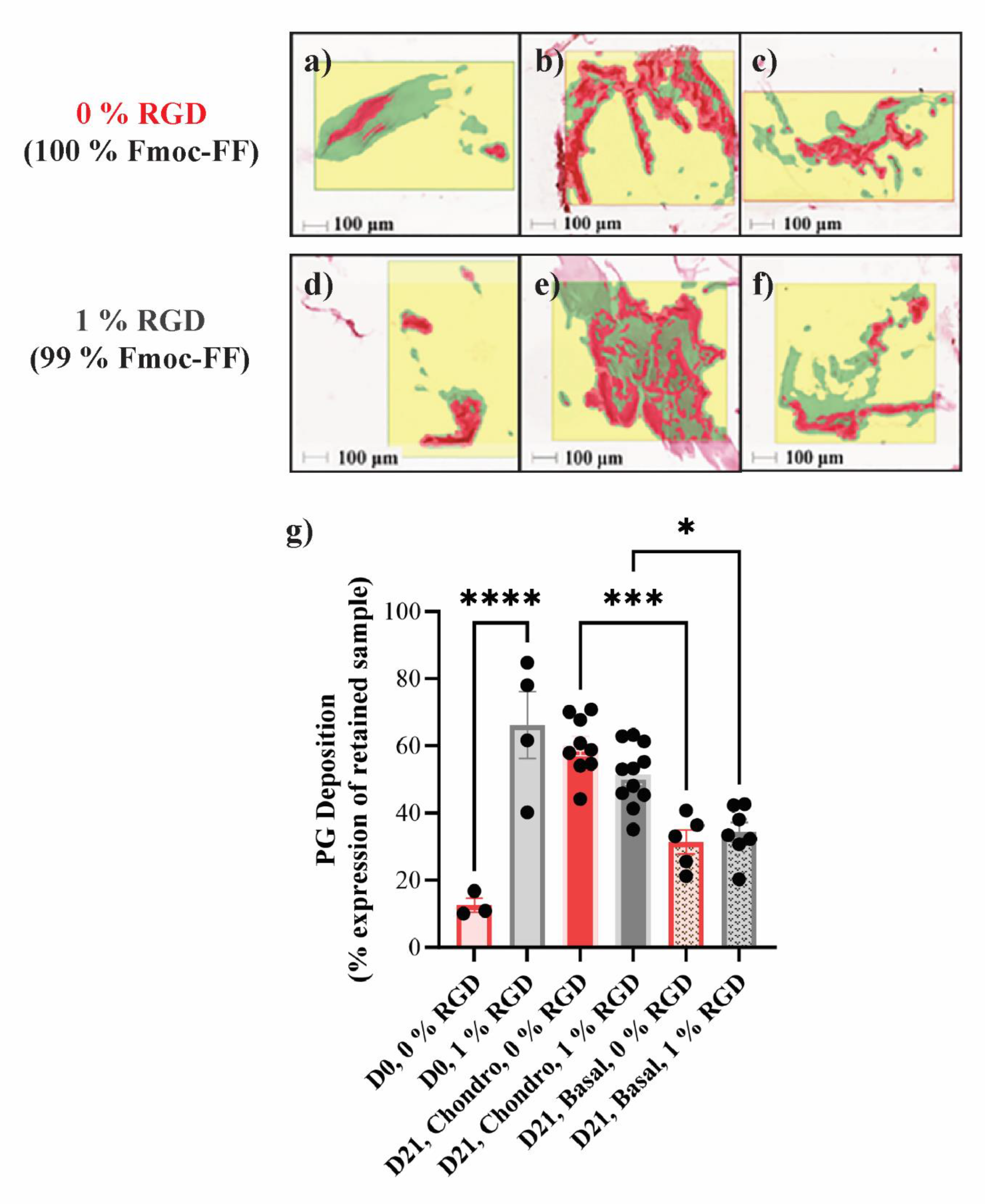
**Safranin-O staining revealed no difference in PG deposition between 0 % and 1 % RGD gels**. Histology to assess and compare proteoglycan (PG) deposition on 0 % RGD, 1 % RGD hydrogels seeded with 250,000 cells per mL. Representative Safranin-O-stained histological slices taken from (**a-c**) 0 % RGD, and (**d-f**) 1 % RGD hydrogels classified for regions of stain expression, retained gel and glass slide features over a 500 µm^2^ section. Quantification of regions of PG expression (**g**) as a percentage of retained Fmoc gel samples paraffinised and sectioned onto glass microscopy slides. • denotes technical replicate. **** = P < 0.0001, *** = P < 0.001, * = P < 0.1.

Quantitative analysis (**figure 5g**) of Safranin O revealed statistically significant greater PG deposition on 0 % RGD and 1 % RGD gels exposed to chondrogenic media than basal media. However, we observed no difference in PG deposition after 21 days between 0 % RGD or 1 % RGD samples (**figure 5g**). The only condition in which we observed much reduced PG deposition was for 0 % RGD prior to incubation in chondrogenic media. We tentatively suggest that this is due to a lack of cell attachment motifs (RGD), limiting PG deposition at short time scales; an effect that is overcome at longer times in culture (day 21, 0 % RGD showed significant PG deposition). However, we cannot rule out issues with sample loss during fixation, and further experiments are required to confirm this hypothesis. This does highlight a need for alternative non-destructive methods to monitor the processes cellular processes occurring *in situ*.

To determine the extent of sGAG deposition by the MSCs, we performed the spectroscopic dimethylmethylene Blue (DMMB) assay. Following papain digestion of the scaffolds, the DMMB dye stains MSC deposited sGAGs in solution and their relative amounts were quantified with UV/Vis absorbance spectroscopy. The results of three independent biological repeats of the DMMB assay are shown in **figure S6**. The results of this data compliment the Safranin O staining and suggest that there is no improvement in GAG deposition in the presence of the RGD ligand after 21 days exposure to chondrogenic media.

### 3.5 Label-free colorimetric histology of collagen reveals the presence of RGD promotes collagen deposition

As type II collagen constitutes over 90 % of native articular hyaline cartilage [64], it is important to develop novel methods to simply and accurately determine the extent of collagen deposition for cartilage tissue engineering applications. Here, we present the first result applying colorimetric histology to collagen imaging [42]. The sensor field of the plasmonic device extends 150 nm from the surface of the device, with material extending beyond this range contributing negligible signal. The typical diameter of the collagen fibres being in the order of 1-10 μm, hence any changes in the colorimetric output are due to differences in the optical properties of the sample (either optical density or birefringence). The goal of the present study was two-fold; first to see if the structure of the collagen fibres could be imaged via colorimetric histology without necessitating any additional staining or labelling, and second, collagen is known to exhibit ordering leading to birefringence [65] which arises due to the ∼63 nm repeating unit in collagen nanofibrils. Hence, assuming an absence of local changes in sample composition we were interested in exploring whether collagen birefringence could be inferred via a colour change on the slide. Quite surprisingly, the deposition of collagen onto the plasmonic devices induced a notable striking colorimetric effect. By adjusting the linear polarisation angle, we were able to dramatically alter the colour contrast on the sample. For the pure collagen hydrogel (positive control), we observe a uniform yellow colour (s-polarisation, **figure 6a**) or uniform red-purple colour (p-polarisation, **figure 6b**) indicating a uniform birefringence throughout the sample.

**Figure 6:**
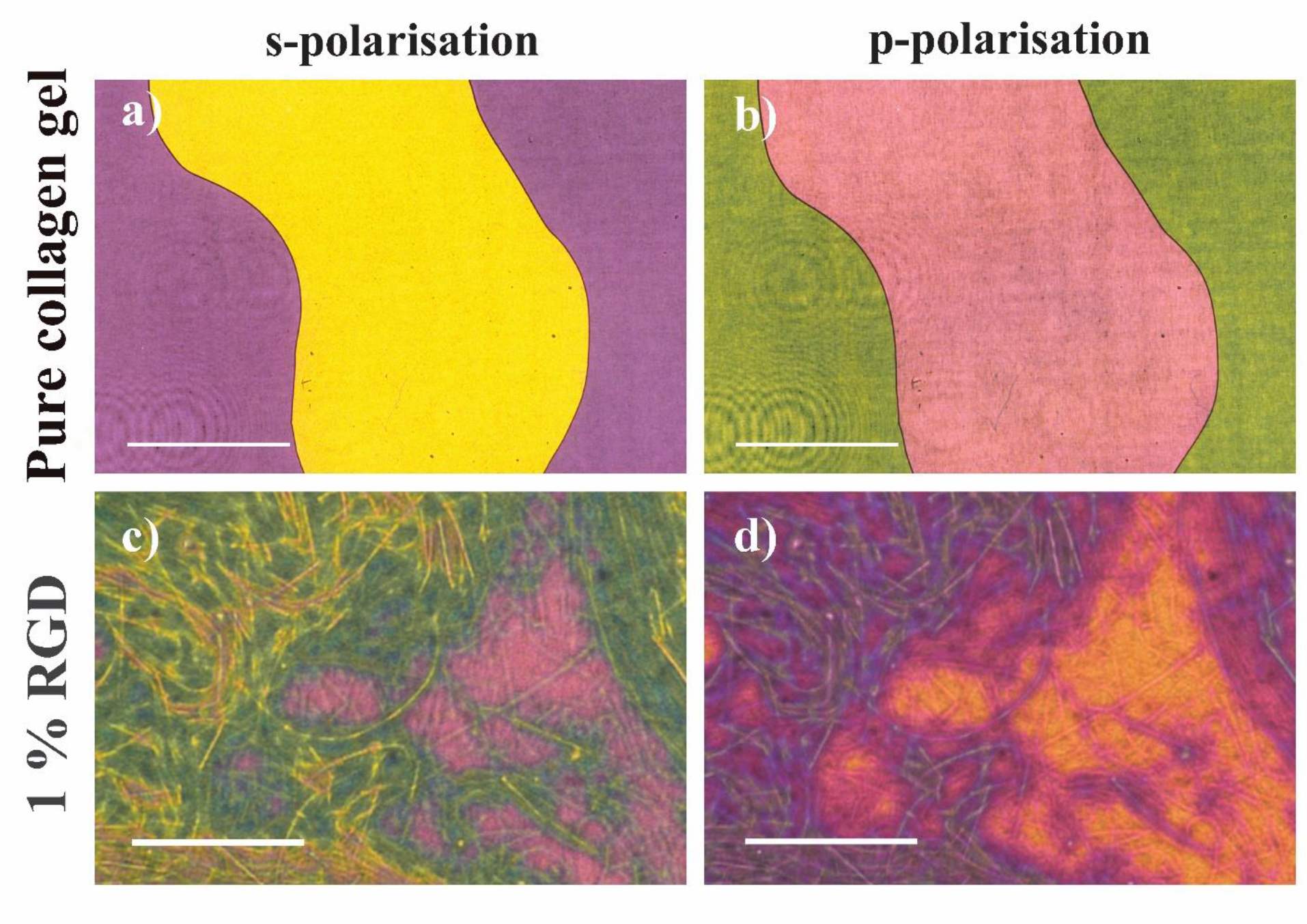
Collagen birefringence is enhanced by colorimetric histology. Birefringence of collagen adsorbed to the colorimetric histology slides is enhanced when viewed through cross-polarised microscopes. Polarised colour changes in transmitted light were observed at **(a)** s-polarisation and **(b)** p-polarisation when pure collagen hydrogels were adsorbed to the slide. Individual bundles of collagen nanofibrils (**c)** and **(d)** could be observed after 21 days culture in chondrogenic media on 1 % RGD gel samples. Images captured at 20x magnification (**a-b**) and 100x magnification (**c-d**). Scale bars **a-b** = 500 µm, **c-d** = 50 µm.

Within MSC-laden Fmoc-FF samples exposed to 21 days in chondrogenic media (**figure 6c-d**), bundles of nanofibrils with sub-micron diameters were clearly observed. Further in many parts of the sample the colour of the sample almost perfectly matched the birefringent collagen-only positive control, with some regions of heterogeneity (e.g. pinker areas in **figure 6c**) suggesting that the fibre bundles may be less ordered in certain regions of the sample. Similar birefringent bundles of collagen fibres were also observed for the other day 21 hydrogels investigated in this study (**figure S7**). These results demonstrate that MSCs cultured within the hydrogels employed in this study exhibited significant collagen deposition. Note it is likely that in the collagen-only samples, the densely packed fibrils within the collagen gels (**figure 6 a-b**), coupled with the microscope’s diffraction-limited resolution, prevent individual collagen bundles from being resolved.

In **figure 7**, we present representative images of applying colorimetric histology to our samples. We observe that there is significant collagen deposition observable after 21 days. No birefringence was observed in the D0 samples (for either the 0 % RGD or 1 % RGD containing samples), which indicated that negligible collagen deposition occurred. In the samples incubated in chondrogenic media for 21 days, we saw clear collagen deposition and birefringence for both the pure Fmoc-FF (0 % RGD) and 1 % RGD containing gels, which indicated that these materials promote collagen deposition. In line with the DMMB and histology results (which showed little difference between samples cultured in chondrogenic or basal media), we also see evidence of collagen deposition for the 0 % and 1 % RGD samples cultured in basal media. We also investigated collagen deposition on gels seeded with higher cell concentrations (500,000 cells), and this revealed very similar trends (**figure S8).** Negative controls composed of 0 % RGD and 1 % RGD containing scaffolds in the absence of cells also revealed no birefringence, further demonstrating that the scaffolds materials are not the source of birefringence (**figure S9**).

**Figure 7:**
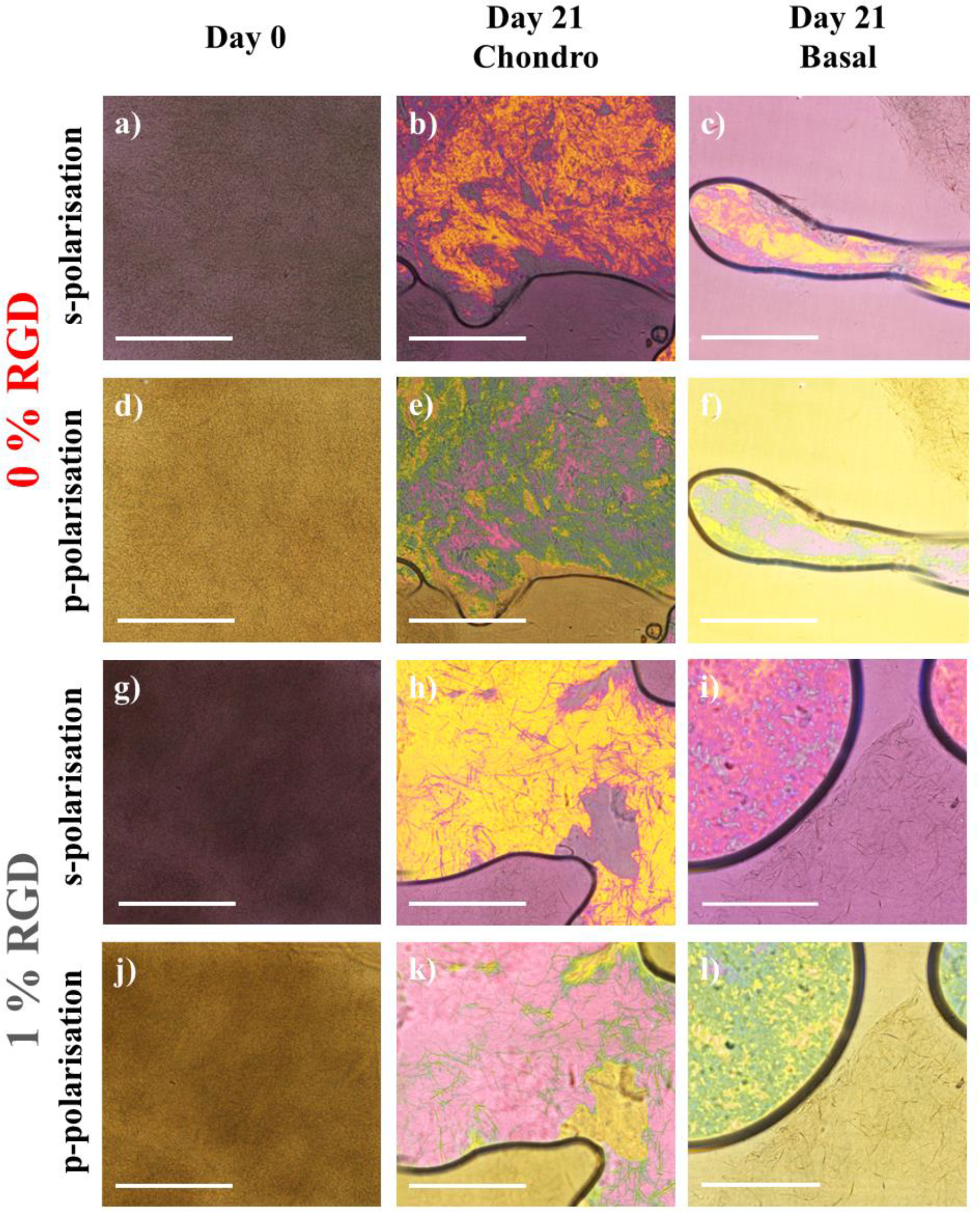
**Extensive collagen deposition is seen after 21 days in culture for all gels**. Plasmon-enhanced colorimetric histology was performed to assess and compare collagen deposition on 0 % RGD and 1 % RGD hydrogels seeded with 250,000 MSCs per mL of gel. (**a-f**) 0 % RGD gels (**g-l**) 1 % RGD samples. (**a-c**) represents s-polarisation (0 % RGD gels) (**d-e**) represents p-polarisation (0 % RGD gels) (**g-i**) represents s-polarisation (1 % RGD gels) and (**j-l**) represents p-polarisation (1 % RGD gels) - 20x magnification, scale bars 500 µm.

We next performed an analysis of the intensity of the birefringence for all the conditions shown in **figure 7**. The stitching capabilities of the Nikon Eclipse Ti2 optical microscope enabled analysis to be performed over an expansive region (each stitched image in **figure S10** is 4.16 x 4.16 mm). Images thresholded against a brightness level determined from the positive control samples and then converted to a hue scale (**figure 8a-b**) previously used to analyse collagen deposition [66, 67] (see methods section 2.11). All regions of gel with hue values greater than the threshold were considered collagen positive, and the mean of the ratio of collagen positive gel to collagen negative gel is plotted in **figure 8c**. We observed statistically significant increases in collagen deposition for the 1 % RGD gels compared to the 0 % RGD gels, which provided convincing evidence that cell attachment (via RGD) to the nanofibrillar network promotes collagen deposition after 21 days in chondrogenic media. Further, as was seen for PG deposition (**figure 5**), the presence of the chondrogenic growth factors BMP-6 and TGF-β in the media promotes collagen deposition, and this has been similarly identified in previous studies of growth-factor inclusive media [68, 69].

**Figure 8:**
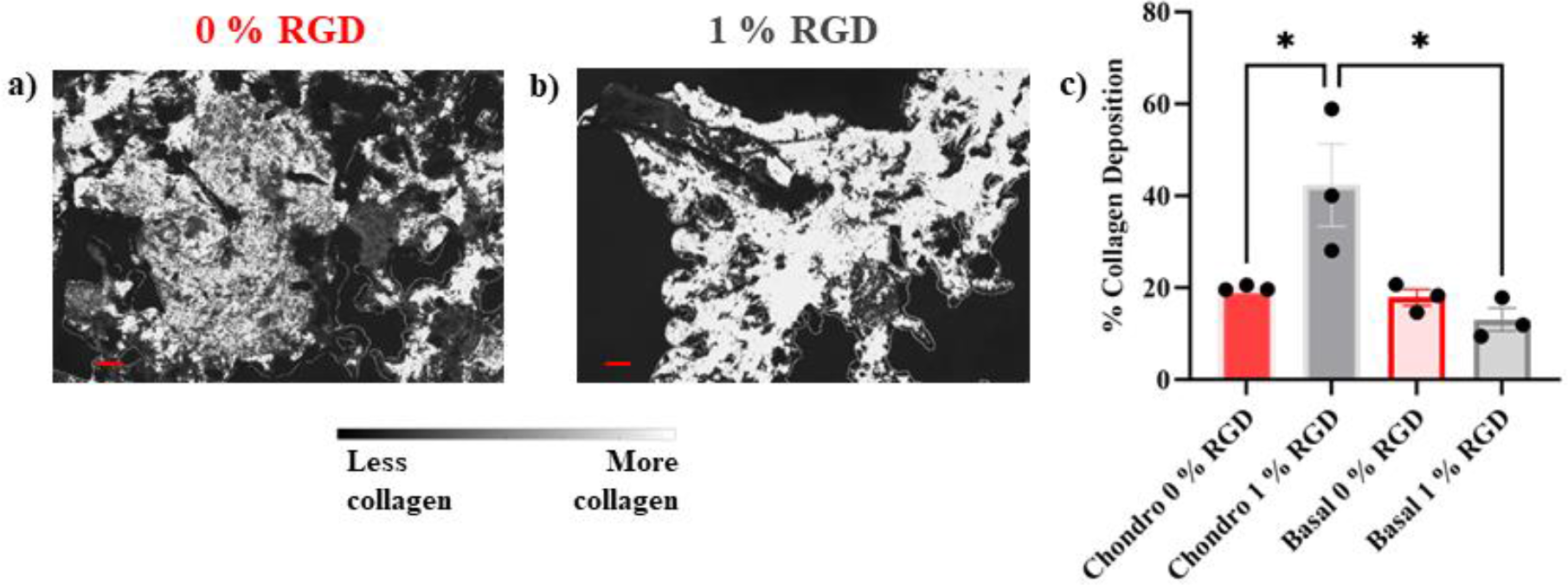
The presence of the RGD ligand significantly increases collagen deposition. Estimate of collagen deposition via colorimetric histology – **a-b)** collagen expression on 0 % and 1 % -RGD gels demarcated based on contrast hues, **c)** quantification of regions of positive collagen deposition (%) in relation to negative collagen deposition for areas of gel deposited on the colorimetric histology slides, quantification performed over at least 3 widefield images per condition. Scale bars = 500 µm. * = P < 0.1

Overall, we observe that MSCs cultured in the Fmoc-FF gels containing 1 % RGD show no increased GAG and PG deposition compared to the pure Fmoc-FF gels. However, the results from the colorimetric histology indicate that the presence of 1 % RGD causes a significant increase in collagen deposition. RGD acts as a cell adhesion moiety by binding to integrins on the cell membrane, thus improving cell adherence to the Fmoc-FF scaffolds [70]. The literature which discusses how cell adhesion affects ECM deposition in MSCs and chondrocytes is often contradictory [71], but Connelly *et. al.* [72] found that RGD-modified hydrogels resulted in reduced sGAG production due to a cell adhesion mediated alteration of the RhoA/Rock pathway. Further, numerous studies have shown that RGD [73, 74] and other cell attachment mediating peptide sequences such as IKVAV from laminin [75], promote collagen deposition of MSCs when encapsulated in hydrogels both *in vitro* and *in vivo*. Thus, we speculate, similar to the studies above, cell adhesion to the RGD functionalised fibrils is promoting collagen deposition perhaps at the expense of PG and sGAG production. The biomolecular mechanisms underpinning this selective improvement in collagen deposition will be the focus of a future study.

## 4. Conclusion

We have shown that the incorporation of RGD into the nanofibrils of self-assembled peptide hydrogels occurs only at low concentrations. By quantifying subtle changes in nanofiber morphology, we were able to show that at higher concentrations Fmoc-RGD no longer incorporates into the fibers and instead acts as a colloidal contaminant preventing stable gelation. Analysis of ECM deposition of MSCs cultured within these hydrogels revealed that the presence of RGD had little effect on sGAG or PG production, however collagen deposition was promoted after 21 days in culture. Further, to overcome the difficulties of *in situ* monitoring of ECM deposition within these gels we report a new application of a recently developed nanofabricated microscope slide for colorimetric histology which allows us to rapidly quantify collagen deposition within biomaterials using nothing more than an optical microscope equipped with a linear polariser. We believe that this label-free, low cost, simple and rapid method of collagen detection will have many applications in a wide range of fields including biomaterials characterisation, disease diagnostics and regenerative medicine.

## Supporting information

Supplemental Information for Chong et al

## Acknowledgements

CC would like to acknowledge the assistance of the students and staff of the La Trobe University Bioimaging Platform, LIMS Histology facility, ACRF Centre for Imaging the Tumour Microenvironment at the Olivia Newton-John Cancer Research Institute, and ARC Centre of Excellence in Advanced Molecular Imaging for their training and assistance with these experiments. NPR would like to acknowledge the La Trobe Institute of Molecular Sciences (LIMS) for the receipt of a Nicholas Hoogenraad fellowship, and Dr. Susi Seibt for assistance on the SAXS / WAXS beamline at the Australian Synchrotron. This research was undertaken, in part, on the SAXS / WAXS beamline at the Australian Synchrotron, part of ANSTO. Figure 1 and the ToC image were partly created using Biorender.

## Table of Contents (ToC) summary and Image

Mesenchymal stem cells cultured in 3D scaffolds can deposit extracellular matrix (ECM) proteins such as collagen, which is beneficial for applications in regenerative medicine. Using a self-assembled peptide scaffold functionalised with cell-adhesive ligands (RGD) it was demonstrated that the addition of RGD promotes collagen deposition, highlighting an innovative nanofabricated plasmonic substrate as a rapid, label-free method of detecting collagen.

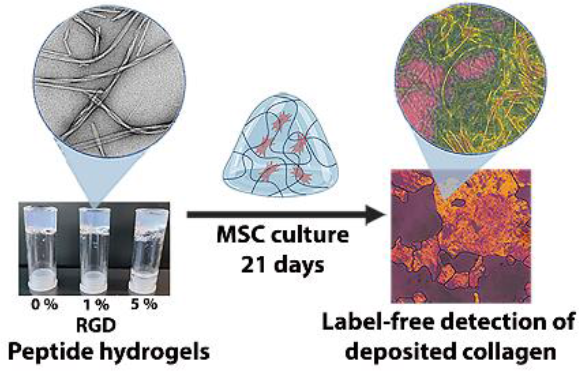

## Data and Conflict of Interest Statements

1. Once accepted all data will be uploaded to a publicly accessible repository (e.g. Zenodo).
2. All authors declare no conflict of interest.

## Notes

### Competing Interest Statement

The authors have declared no competing interest.

### Summary of Updates

Manuscript revised to include data on ECM deposition and colorimetric histology

