## Supplemental Information for Chong et al for "Extensive collagen deposition by mesenchymal stem cells cultured in 3D self-assembled peptide scaffolds as revealed by nanoplasmonic colorimetric histology"

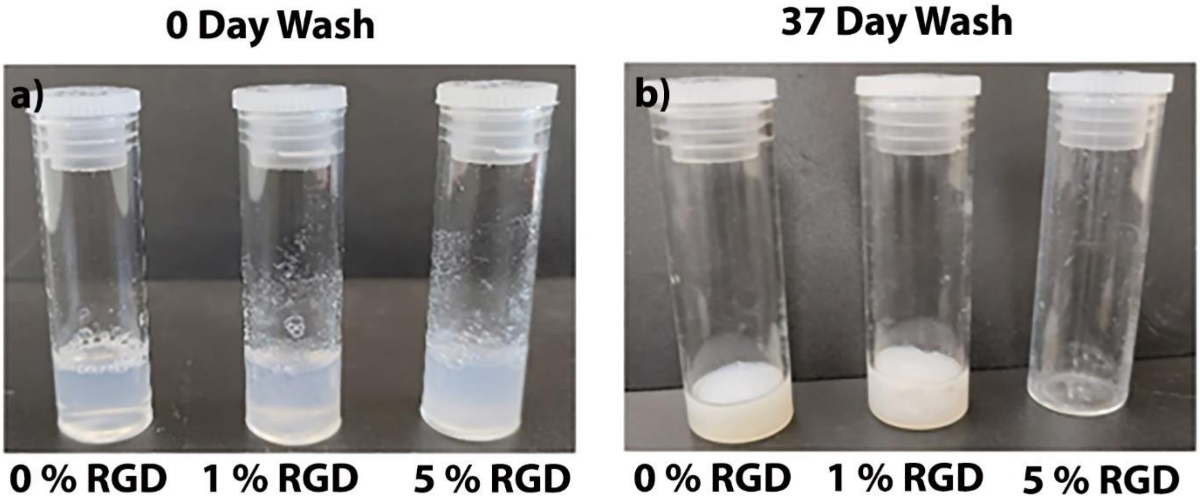


**Figure S1**: **Degradation hydrogels pre and post periodic DMEM (minus phenol red) wash steps. a)** 0 % RGD, 1 % RGD, and 5 % RGD gels pre-commencement of wash steps and on completion of daily wash steps in DMEM for 37 days.

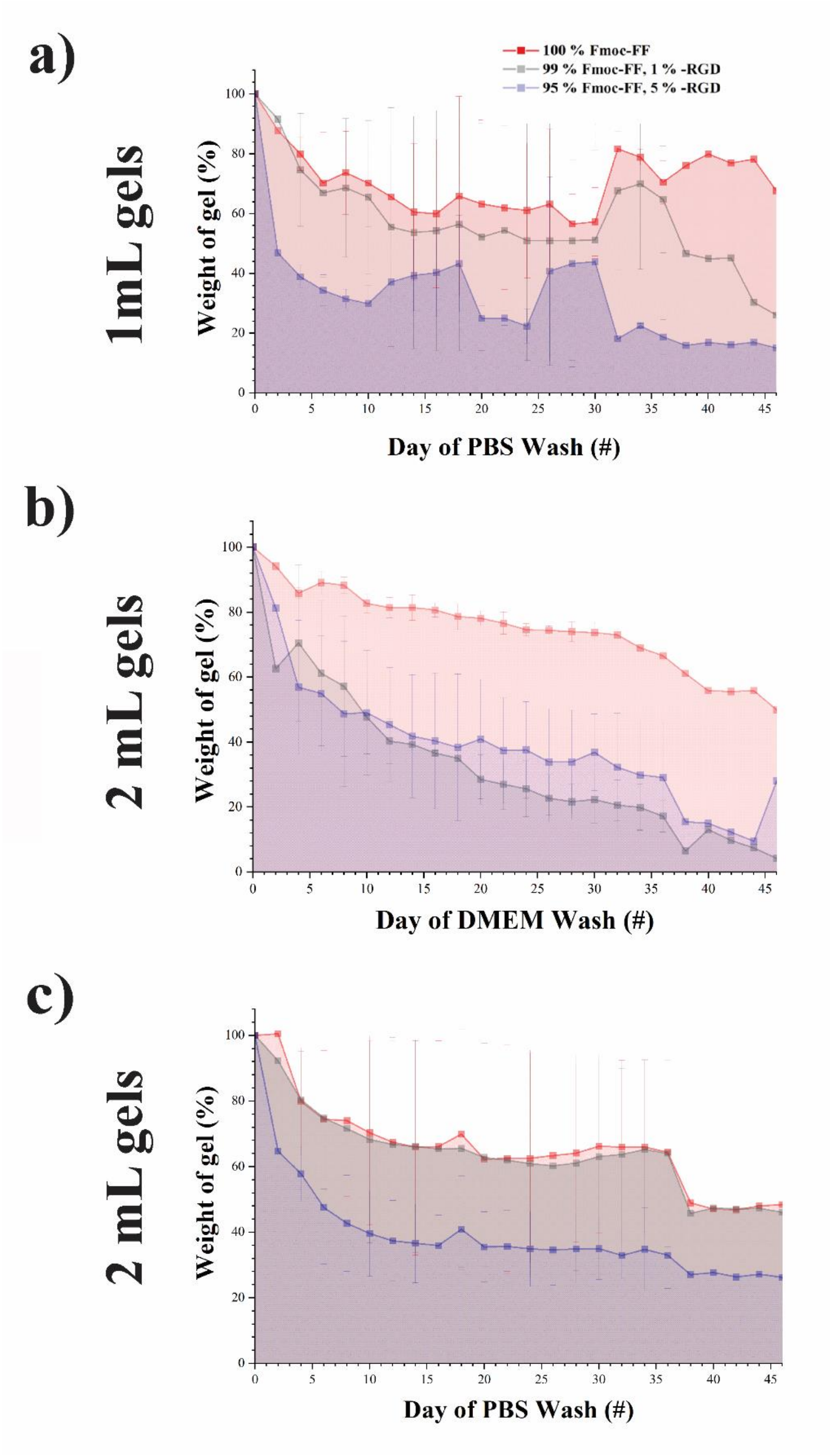


**Figure S2**: **Rate of gel degradation is affected by RGD concentration.** Mass loss assays for 0 % RGD, 1 % RGD and 5 % RGD hydrogels. **a)** 1 mL hydrogels in PBS. **b)** 2 mL hydrogels in DMEM. **c)** 2 mL hydrogels in PBS. Error bars denote mean ± SD for n = 2.

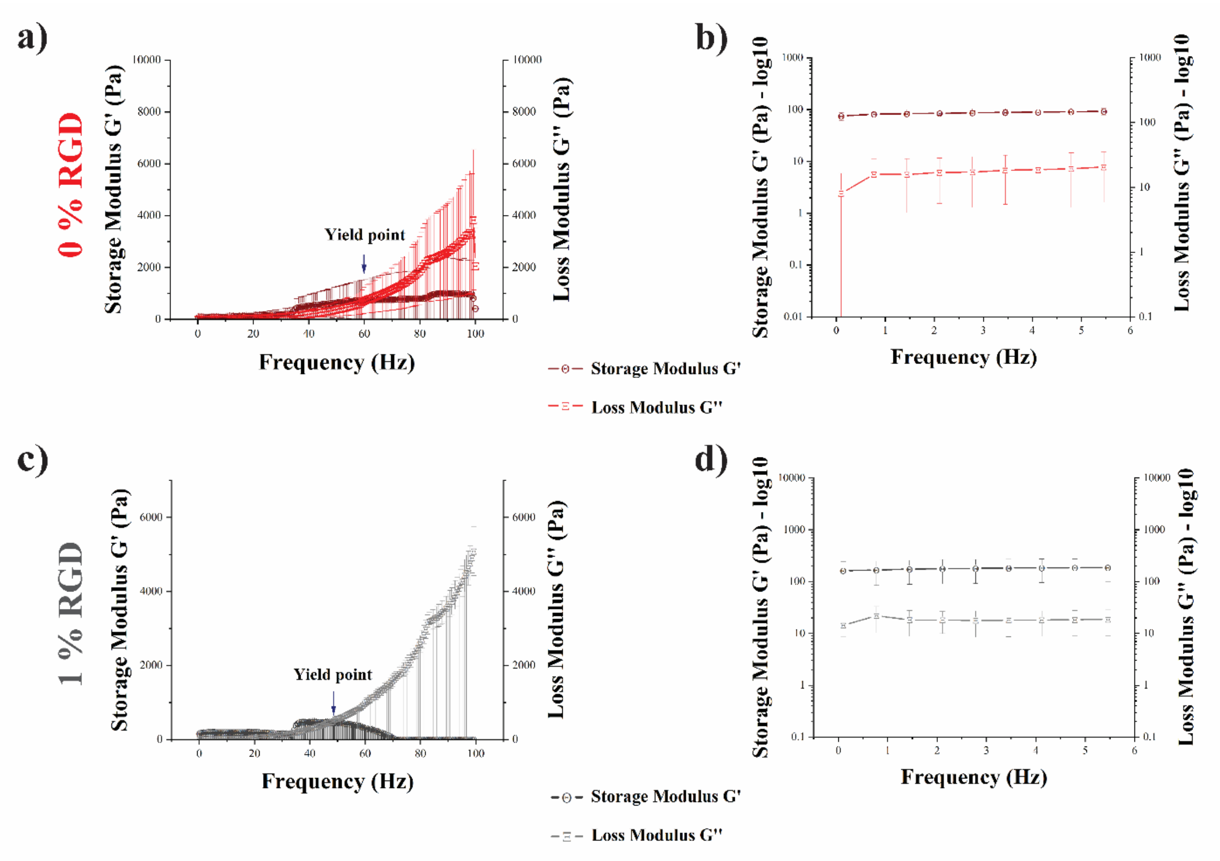


**Figure S3**: **Additional rheological data shows gel breakage at high frequency of oscillation. a-b)** 0 % RGD hydrogels, **c-d)** 1 % RGD hydrogels. At frequencies of 60 Hz and 50 Hz respectively, G" (loss modulus) exceeds G' (storage modulus), marking a yield point, and a transition point of the gels into a liquid state. These gels may thus be prone to collapse under shear force.

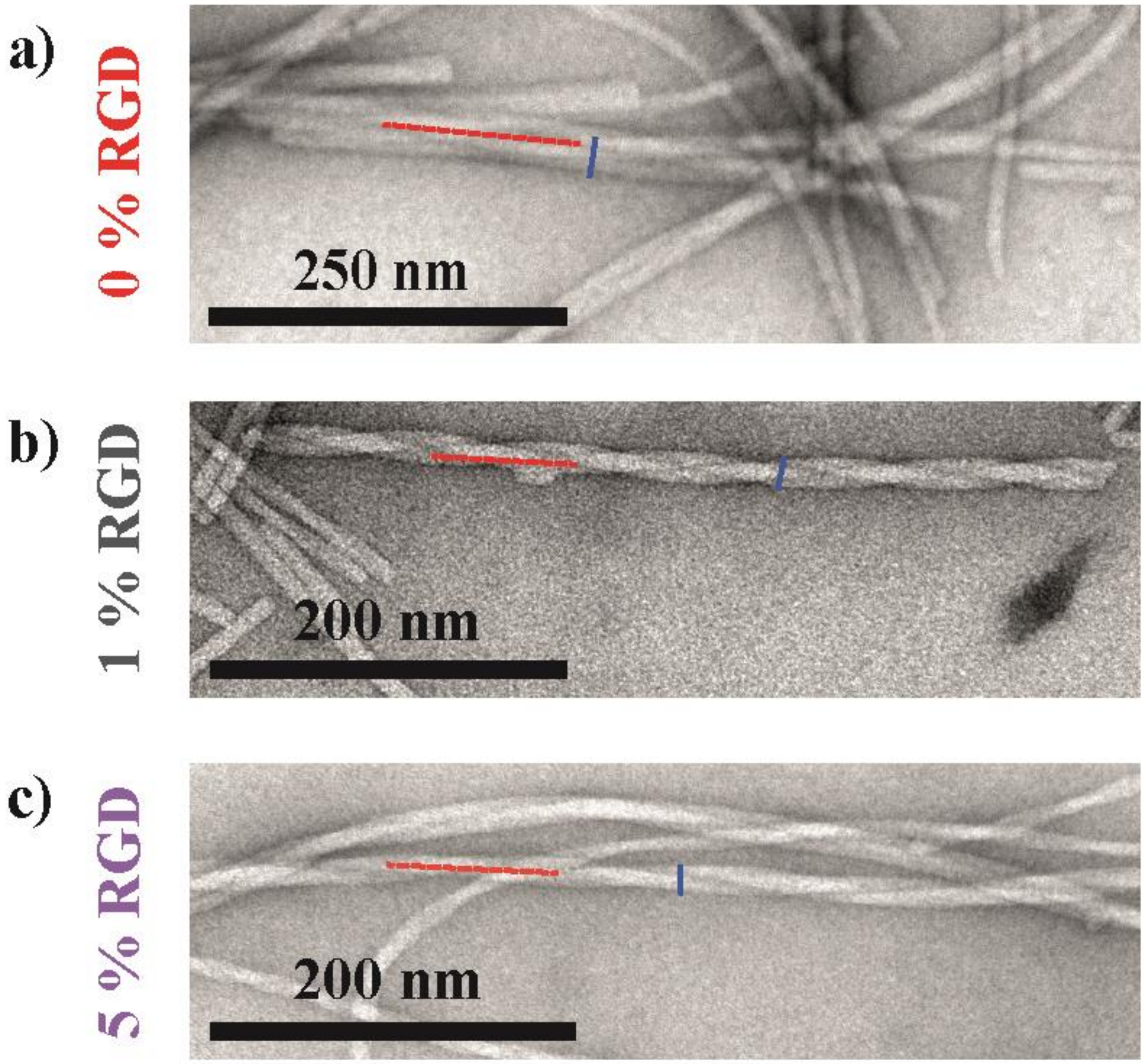


**Figure S4**: **Fibrillar helical periodicity was observed across TEM images of respective hydrogels. a)** 0 % RGD hydrogels. **b)** 1 % RGD hydrogels. **c)** 5 % RGD hydrogels. Peak-to-peak pitch (red broken line), and fiber width pitch (blue solid line) is indicated within each helical ribbon.

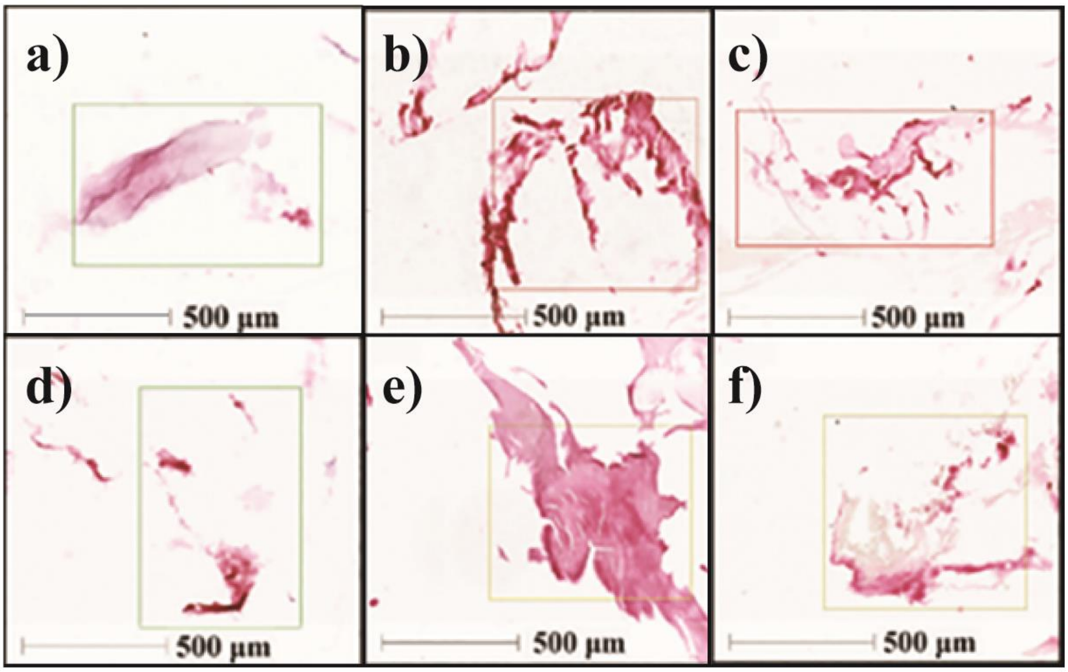


**Figure S5: Safranin-O staining revealed no difference in PG deposition between 0 % and 1 % RGD gels**. Histology to assess and compare proteoglycan (PG) deposition on 0 % RGD, 1 % RGD hydrogels seeded with 250,000 cells per mL. Representative Safranin-O-stained histological slices taken from (**a-c**) 0 % RGD, and (**d-f**) 1 % RGD hydrogels. Selected regions of 500 µm$^2$ were selected from each sample set, and slide features were then demarcated and quantified.

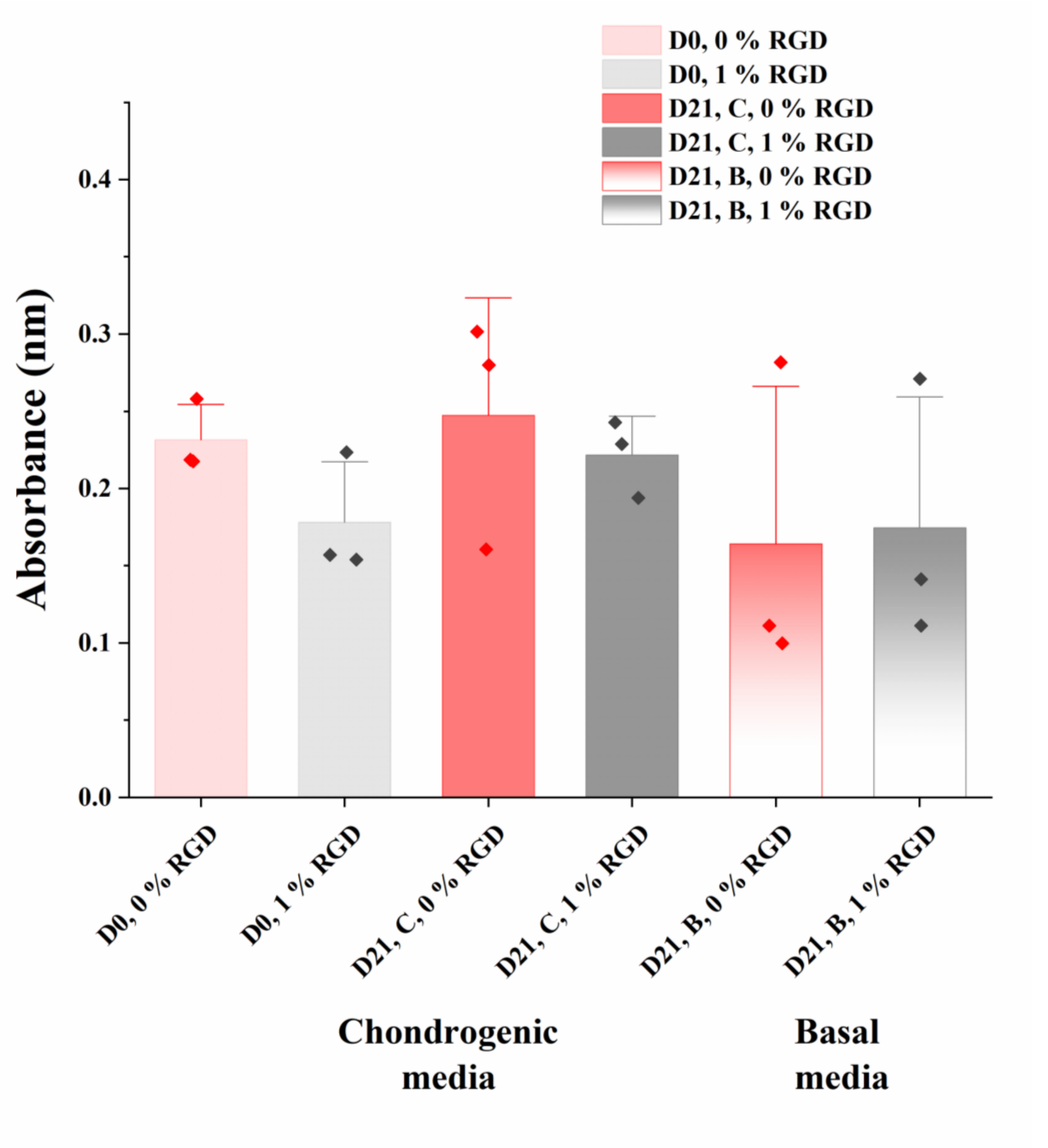


**Figure S6: Dimethylmethylene Blue (DMMB) assay reveals no significant changes in sulphated glycosaminoglycans (sGAGs) deposition.** Absorbance readings were obtained at 595 nm for 0 % RGD and 1 % RGD samples at days 0 and 21 respectively. C = chondrogenic media, B = basal media. All samples seeded with 250K cells per mL and exposed to chondrogenic or basal media up to 21 days.

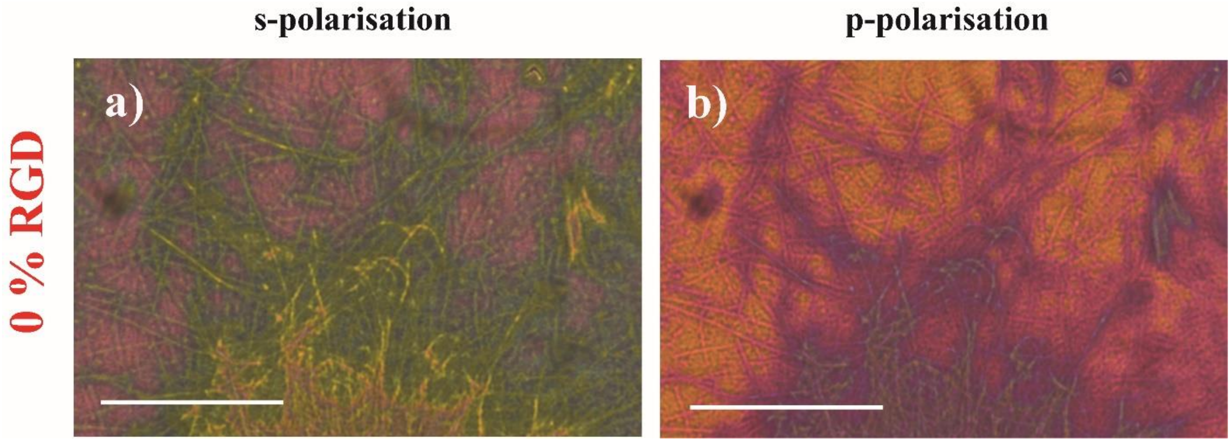


**Figure S7: Collagen deposition observed on 0 % RGD containing hydrogels (seeded with 500,000 cells) using high magnification (x100).** Clear evidence of birefringent collagen bundles can be seen after 21 days culture in chondrogenic media (scale bars = 100 µm).

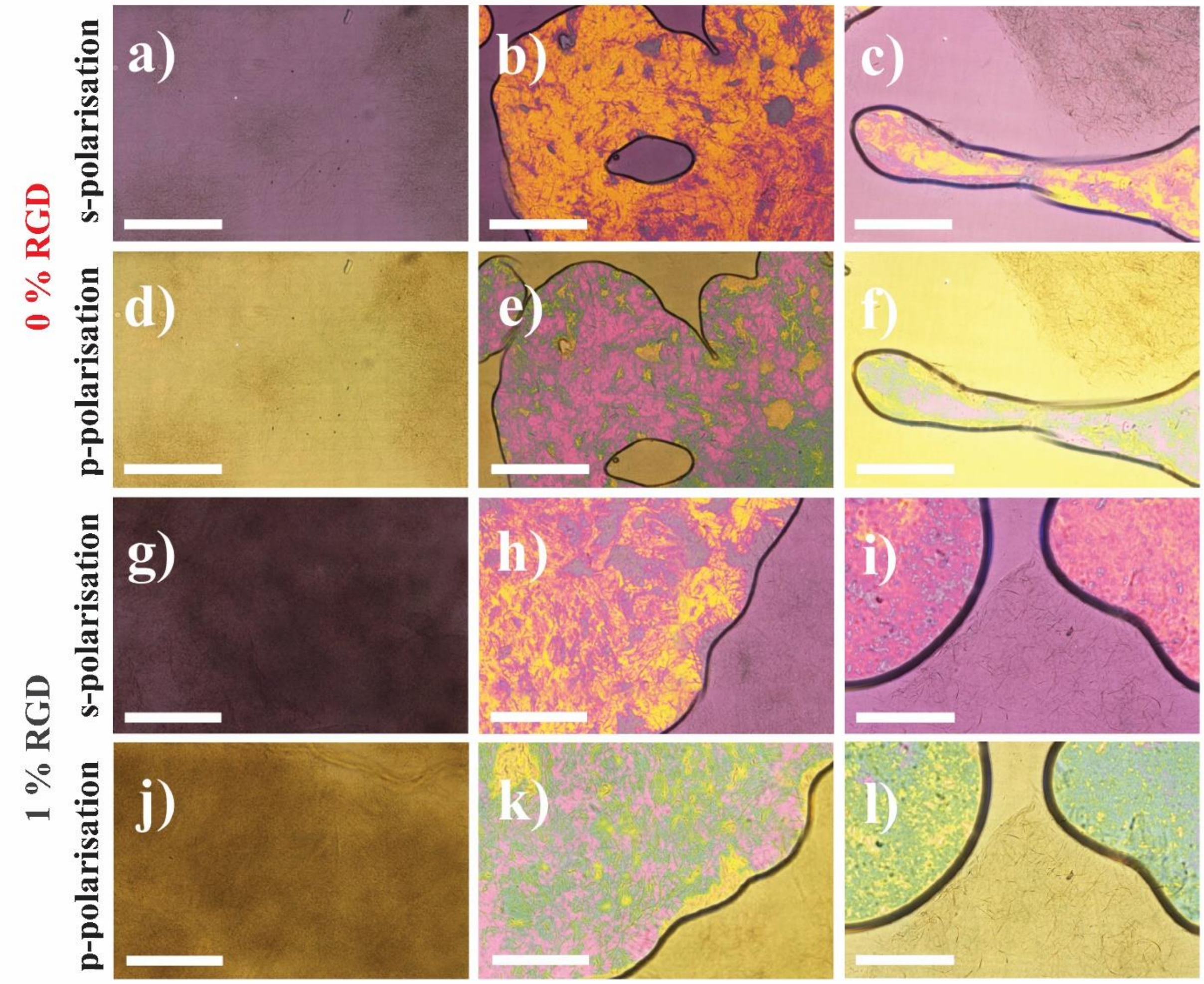


**Figure S8: Nanoplasmonic colorimetric histology also shows extensive collagen deposition for MSCs cultured within 0 % RGD and 1 % RGD hydrogels when seeded at 500,000 MSCs per mL**. Polarised colour changes in transmitted light were observed at s-polarisation (**a-c, g-i**) and p-polarisation (**d-f, j-l**) resonances. Cell-seeded samples were initially imaged at day 0. Samples were then exposed to chondrogenic or basal media, and imaged at day 21 post-cell seeding (scale bars = 500 µm).

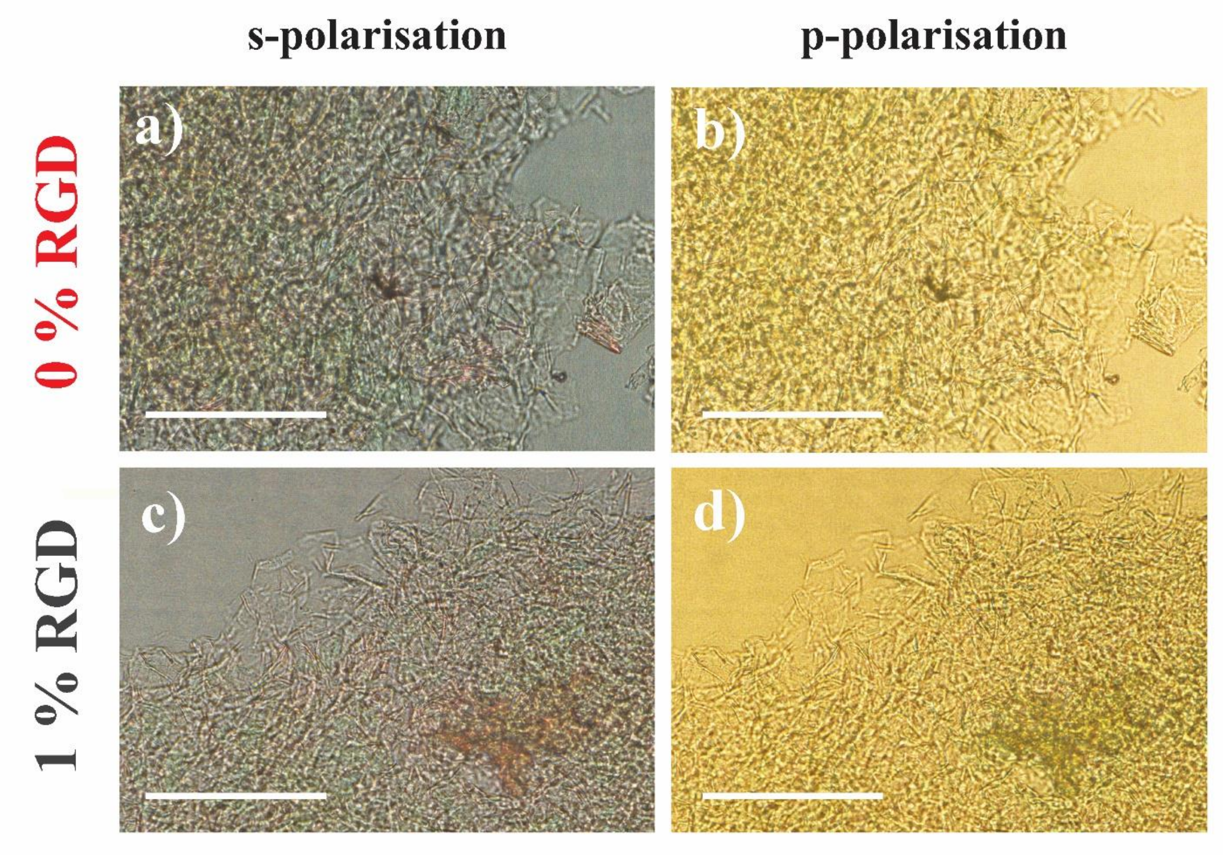


**Figure S9: Plasmonic colorimetric histology negative controls showed no birefringence.** 0 % RGD without cells (**a-b**) and 1 % RGD without cells (**c-d**) (scale bars = 500 µm).

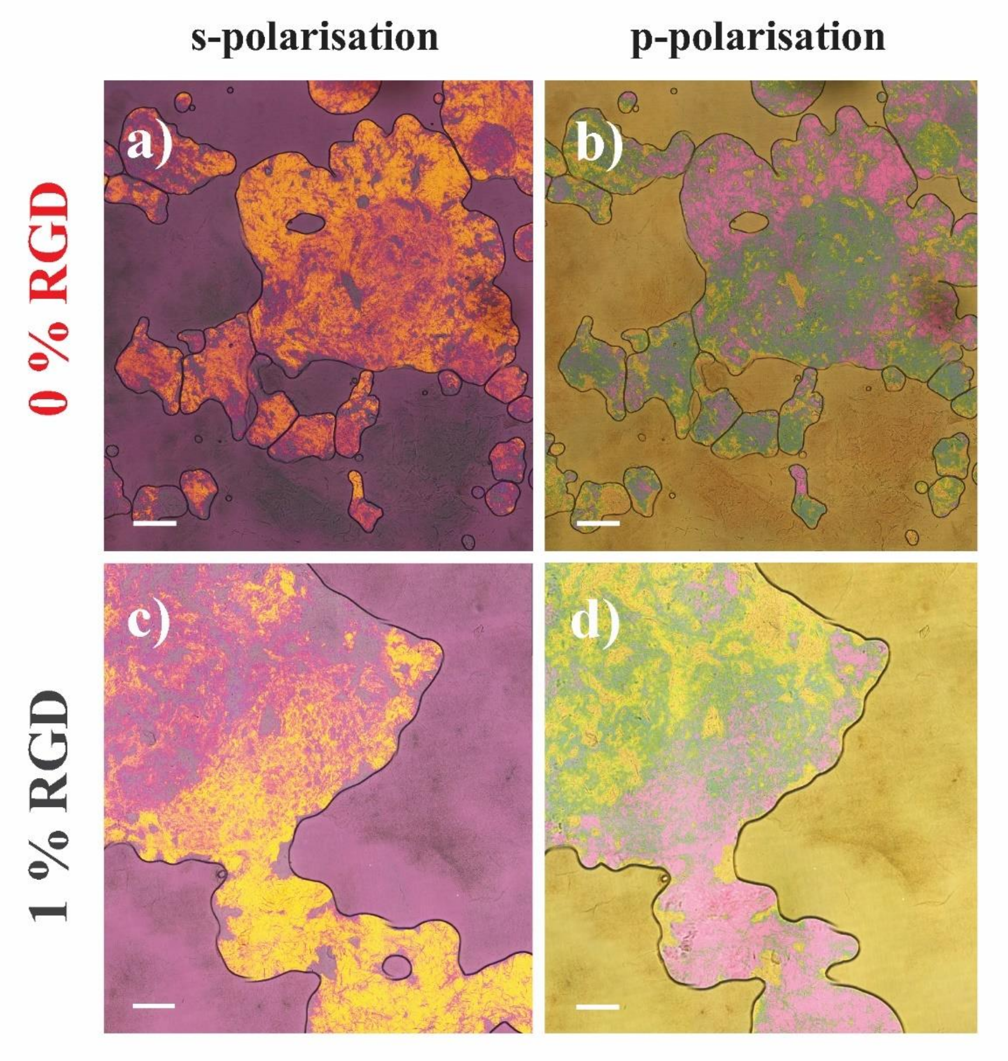


**Figure S10: Widefield comparison of birefringence / collagen deposition on 0 % RGD and 1 % RGD gels at day 21.** Polarised colour changes in transmitted light were observed on 0 % RGD cell-seeded gels at s-polarisation (**a**) and p-polarisation (**b**), and 1 % RGD cell-seeded gels at s-polarisation (**c**) and p-polarisation (**d**). Samples were initially seeded with 250,000 cells each at day 0, and then imaged at day 21 (scale bars = 500 µm). Image stitching was performed with NIS Elements AR software (Nikon, Tokyo, Japan) with each image representing a 4.16 x 4.16 mm field.
